# Enhancer-promoter interactions are reconfigured through the formation of long-range multiway chromatin hubs as mouse ES cells exit pluripotency

**DOI:** 10.1101/2023.02.08.527615

**Authors:** D. Lando, X. Ma, Y. Cao, A. Jartseva, T. J. Stevens, W. Boucher, N. Reynolds, B. Montibus, D. Hall, A Lackner, R. Ragheb, M Leeb, B. D. Hendrich, E. D. Laue

## Abstract

Enhancers are genomic DNA sequences that bind transcription factors, chromatin regulators and non-coding transcripts to modulate the expression of target genes. They have been found to act from different locations relative to a gene and to modulate the activity of promoters up to ∼1 Mb away. Here we report the first 3D genome structures of single mouse ES cells as they are induced to exit pluripotency, transition through a formative stage and undergo neuroectodermal differentiation. In order to directly study how interactions between enhancers and promoters are reconfigured genome wide we have determined 3D structures of haploid cells using single cell Hi-C, where we can unambiguously map the contacts to particular chromosomes. We find that there is a remarkable reorganisation of 3D genome structure where inter-chromosomal intermingling increases dramatically in the formative state. This intermingling is associated with the formation of a large number of multiway hubs that bring together enhancers and promoters with similar chromatin states from typically 5-8 distant chromosomal sites that are often separated by many Mb from each other. Genes important for pluripotency exit establish contacts with emerging enhancers within multiway chromatin hubs as cells enter the formative state, consistent with these structural changes playing an important role in establishing new cell identities. Furthermore, we find that different multiway chromatin hubs, and thus different enhancer-promoter interactions, are formed in different individual cells. Our results suggest that studying genome structure in single cells will be important to identify key changes in enhancer-promoter interactions that occur as cells undergo developmental transitions.

## Introduction

Embryonic stem (ES) cells can differentiate into all the different lineages of a mature organism. Naïve and primed ES cells *in vitro* correspond to the pre- and post-implantation populations in the embryo^1,2^, and they are developmentally linked to each other via progression through a ‘*formative’* transition state where cells mature in response to inductive cues (Figure 1A)^3^. In this formative state, there is a rewiring of transcription factor networks – this results in the downregulation of genes required for naïve pluripotency, such as *Nanog and Klf4,* and the upregulation of genes associated with the primed epiblast, such as *Otx2* and those encoding the de novo methyltransferases *Dnmt3a/b*. After transition through the formative state, epiblast cells can be induced to form primordial germ cells and subsequently they become progressively specified towards different cell fates^4-7^. Studying the formative state thus allows us to understand the mechanisms of multi- lineage decision-making. Stimulated by the observation that nuclei of formative state cells have different material properties to those of either naïve or primed ES cells^8^ we have explored how genome structure changes in this crucial transition state.

**Figure 1.**
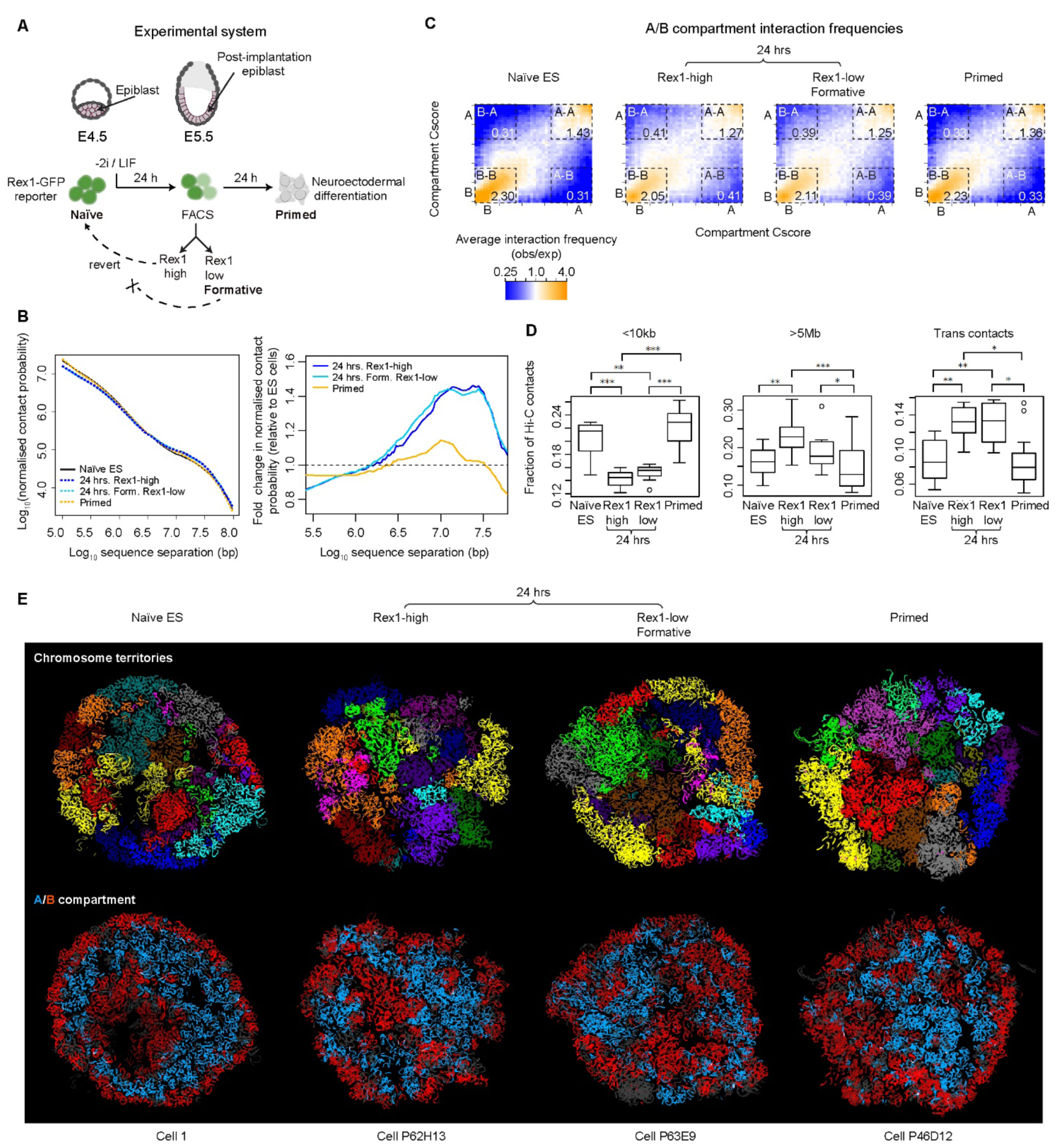
The 3D genome of mouse ES cells goes through a structural transition in the formative state. **A)** Twenty-four hours after two inhibitors (of the MEK/Erk pathway and glycogen synthase kinase-3) and Leukaemia inhibitory factor (2i/LIF) are withdrawn the Rex1 (*Zfp42*) gene becomes silenced as cells exit naïve pluripotency: at this time a proportion of cells have progressed into the formative (Rex1-low) state, whilst the remainder (Rex1-high cells) are still exiting naïve pluripotency^4^. A further 24 hrs later, as *Zfp42* expression is lost in all cells, mouse ES cells enter a primed state that in basal media prepares them to differentiate down the neuroectodermal lineage. **B)** (Left) plots of log10 contact probability against log10 intra-chromosomal sequence separation determined using in-nucleus Hi-C experiments on large populations of cells. (Right) expansion showing the fold change in contact probability relative to that in naïve ES cells. **C)** Saddle plots showing the normalised average A/B compartment interaction frequencies between pairs of 25 kb bins, arranged by their A or B compartment Cscore’s. The numbers in the corners show the average normalised Cscore within the boxed A/B compartment regions (see the Methods for the definition of the Cscore). **D)** Boxplots showing the fraction of contacts in the single nuclei Hi-C datasets that are: <10 kb apart in the same chromosome (left); >5 Mb apart in the same chromosome (middle); or between different chromosomes (right). The midline is the median value, the lower and upper edges of the box represent the 1st and the 3rd quartiles, and the whiskers represent the minimum and maximum values. False Discovery Rate (FDR) adjusted Mann Whitney U test *p*-values are also shown: * - *p* < 0.05; ** - *p* < 0.01; *** - *p* < 0.001. **E)** 3D genome structures of representative single G1 phase nuclei, where the individual chromosomes are either coloured differently (top), or coloured blue or red depending on whether a particular region of a chromosome is in the A or B compartment, respectively (bottom).

3D genome structure is closely linked to many biological processes in the nucleus including transcription, DNA replication and DNA repair – its alteration is also strongly implicated in developmental and degenerative disease. In recent years a very large number of experiments employing light microscopy and either biochemical ligation-based (chromatin condensation capture - 3C) or ligation-independent methods (split-pool recognition of interactions by tag extension - SPRITE; genome architecture mapping - GAM), have suggested that three key mechanisms – compartmentalisation, loop extrusion and tethering – play a critical role in organising 3D genome structure in the nucleus. Biochemical experiments carried out on populations of cells using the 3C, SPRITE and GAM methods have illuminated important features of chromosome and genome structure that are conserved in many (or all) cells. In particular, they have shown that the genome is segregated into: *1)* A and B compartments (eu- and hetero-chromatin); *2)* megabase-scale topologically associating domains (TADs), which have a higher frequency of intra- compared to inter-domain chromatin interactions; and *3)* loops where specific regions of the genome contact each other more frequently. Both TADs and loops are generated via loop extrusion mechanisms leading to convergent CTCF sites becoming linked through bridging interactions via Cohesin. Importantly, evidence suggests that enhancers can be bought into close physical proximity with their target promoter by this process. Alongside these studies a combination of imaging and sequencing based methods have shown that Lamin-associated domains are either tethered to the nuclear periphery (to form LADs) or sequestered around the nucleolus (to form NADs). For recent reviews of the biochemical techniques used to study genome organization, the mechanisms of chromosome folding and nuclear organization that have been uncovered, and the mechanisms of enhancer action, see Refs. ^9^, ^10^ and ^11^, respectively.

The extraordinary variability of chromosome/genome structure from cell-to-cell means that single cell studies are needed for 3D structure determination^12-18^. Many structural features, such as the intermingling between chromosomes and the 3D positions of individual enhancers and genes, are hidden in biochemical experiments carried out on populations of cells because they measure a statistical average of contact probability across many (often millions) of cells. By contrast single cell Hi-C experiments (one of the 3C methods) allow one to unambiguously map individual pairwise contacts and therefore study particular enhancer-promoter interactions. In addition, the complete set of contact data can be used as constraints to calculate 3D structures of how chromosomes/intact genomes fold in single cells^12-14,17^. 3D structure determination is also useful because it allows one to distinguish real long-range contacts from noise in single cell Hi-C experiments.

Moreover, it is possible to determine the relative 3D positions of genes that are far away from each other in the nucleus, and which don’t contact each other^19^. The enormous variability of chromosome and genome structure from cell-to-cell that has been observed in single cell Hi-C experiments has been confirmed by recent advances combining highly multiplexed fluorescence in-situ hybridisation (FISH) and super-resolution imaging which allows the tracing of chromatin structure in single cells. Either individual enhancer-promoter interactions can be imaged at high-resolution, or the organisation of e.g. TADs within a chromosome can be studied at a lower resolution. In further developments of this technology, chromatin tracing has been combined with studies of RNA levels of hundreds of genes. For a review of the large number of exciting new methods that have recently been developed to image the 3D genome and trace chromatin structure, see Ref.^20^.

Previous studies have shown that there are significant changes in chromosome structure during the parental to embryo switch as gene expression is first activated^15,21,22^. In addition, changes in chromosome structure that correlate with changes in transcription can be observed over days and weeks in longer-term development^23-25^. However, it has been unclear how enhancer-promoter interactions change genome-wide as cells undergo rapid alterations in the expression of large numbers of genes during cell fate transitions (over 1-2 cell cycles). In this work we wanted to study how 3D genome structure changes as cells exit pluripotency, and to carry out an unbiased genome-wide study of enhancer-promoter interactions in single mouse ES cells in the naïve, formative and primed states of pluripotency. Unexpectedly, we find that there is a major reorganisation of 3D genome structure as cells transition from the naïve into the formative pluripotent state. We find that promoters of genes whose expression levels are changing during this transition become co-located with enhancers that reside in sequences that are often >10 Mb distant in structures we call “multiway chromatin hubs”. Importantly, the multiway chromatin hubs formed, and the identities of the long-range interacting sequences involved, varies from cell to cell. These multiway chromatin hubs, which cannot be detected using experiments in which populations of cells are studied, are located at the interface between the part of the chromosome that is buried and the regions that become highly intermingled with other chromosomes. They are dependent upon the DNA methylation machinery for their formation, and they correlate with areas of Polycomb repressive complex activity. By determining 3D genome structures in single cells, we provide evidence that promoters of genes required for pluripotency exit establish unusually long-range interactions with different emerging enhancers in different individual cells to drive the transcriptional changes necessary for lineage commitment.

## Results

### Mouse ES cells undergo a structural transition shortly after the onset of differentiation

We exploited an *in-vitro* system which recapitulates the early *in vivo* stages of mouse ES cell differentiation down the neuroectodermal lineage^4,7^ to ask whether genome structure changes as cells exit from pluripotency into the formative state (Figure 1A). In-nucleus Hi-C experiments (Supplementary Table 1) using mouse ES cells expressing a Rex1-destablised GFP fusion^26^ revealed that 24 hrs after induction of differentiation the genome of mouse ES cells goes through a distinct structural transition in the Rex1 (*Zfp42*)-high (less differentiated) and low (formative) states, with a decrease in short and an increase in long range intra-chromosomal Hi-C contacts (<1 Mb and >5 Mb, respectively, Figure 1B). Twenty-four hours later Hi-C contact probability reverts to naïve ES-like levels as cells progress towards the primed state (Figure 1B). We found that formation of the 24 hr transition state involves a weakening in A/B compartment insulation (Figure 1C) – i.e. in interactions between euchromatin and heterochromatin^10^.

To understand these structural changes we carried out single nucleus (sn) Hi-C experiments to determine 3D genome structures^12,14^ (Supplementary Table 2). In both Rex1-high and low cells we observed an increase in inter-chromosomal contacts in the 24 hr state as well as a loss of short- and increase of long-range (>5 Mb) intra-chromosomal contacts (Figure 1D). Calculation of 3D genome structures of individual G1 phase nuclei (Supplementary Table 3) showed that both naïve ES and primed cells have a characteristic Rabl configuration of chromosomes, but this is very disrupted in the 24 hr state (Supplementary Figure 1). Mapping of the A/B compartments on to the 3D single cell structures showed that the naïve ES cell structure – which comprises a ring of B compartment at the nuclear periphery and around the nucleolus sandwiching the A compartment in the interior^14^ – becomes fragmented in the 24 hr state (Figure 1E). In primed cells A/B compartment structure partially reverts towards that seen in the naïve ES cell state. Strikingly, however, in both Rex1-high and low cells, we found that the structural disruption in the 24 hr state mainly involves a very significant increase in intermingling between different chromosomes in the A-compartment compared to naïve ES and primed cells (Figures 2A,B; see also Supplementary Videos 1–4). The shorter cell cycle in formative state cells^27^ suggested that this greater intermingling does not result from chromosomes having more time to unfold in G1 following mitosis.

**Figure 2.**
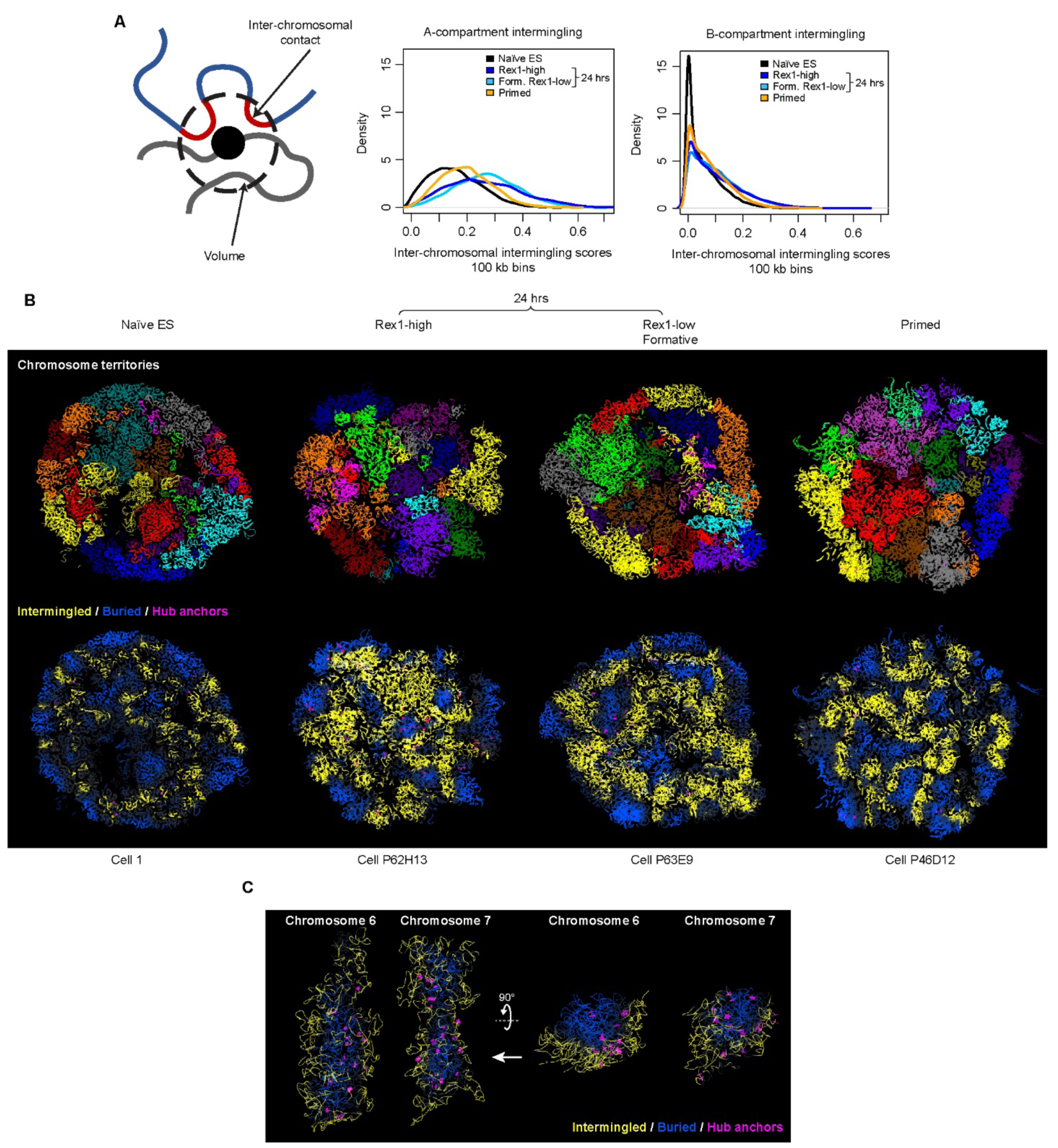
The structural transition in the formative state involves increased chromosomal intermingling in the A-compartment. **A)** (Left) An intermingling score was defined as the fraction of beads (in the polymer model of the 3D genome structure) that are located on different chromosomes and in close spatial proximity. (Right) The inter-chromosome intermingling scores of genomic loci determined from the 3D structures, and plotted as density curves, shows a significant increase in the A compartment in 24 hr state cells (on average 63% higher in Rex1-high and 81% higher in Rex1-low (formative) vs naïve ES cells, Mann Whitney U test for both cases: *p* < 10^-15^). (Note – there is also a statistically significant, but much smaller increase in intermingling in the B compartment in 24 hr cells.) **B)** Slices through 3D genome structures of representative single G1 phase nuclei, where the individual chromosomes are either coloured differently (top), or coloured yellow or blue depending on whether that particular region of the chromosome is either intermingled with another chromosome or buried (bottom). (Regions transitioning between the two are shown in grey.) **C)** Exploded views of part of the 3D structure of a Rex1-low, formative state cell showing the isolated chromosomes 6 & 7 in two different orientations (left & right). The chromosomes are coloured as in B) and the positions of the ‘anchors’ of the chromatin hubs are indicated by magenta stars. [In B), five independently calculated 25 kb structures are superimposed to show that the structures are well determined by the Hi-C data, whilst in C) a single 100 kb structure is shown so that the positions of the hub anchors can be seen more clearly.]

Previous work carried out on populations of cells has shown that ‘*stripes*’ of Hi-C contacts arise from the extrusion of DNA loops by Cohesin^28-30^. Hi-C and higher resolution Micro-C data additionally revealed the presence of ‘*dots*’ within such stripes^31-33^. In our single nucleus Hi-C experiments of 24 hr state cells we were often able to directly observe a similar series of Hi-C contacts aligned along the same horizontal or vertical axis in the contact maps (Figure 3A). However, because these series of contacts all occur in the same single haploid nucleus, and cannot arise from contacts found in different cells, the snHi-C experiments show that they correspond to ‘*multiway chromatin hubs*’ where genomic interactions form between an ‘*anchor*’ (a single small ∼40 kb sequence) and multiple (typically 5-8) *‘contact*’ sites. Generation of 3D structures from the Hi-C data allowed us to study the spatial arrangement of these hubs^19^ that are often far apart from each other in the nucleus (see below).

**Figure 3.**
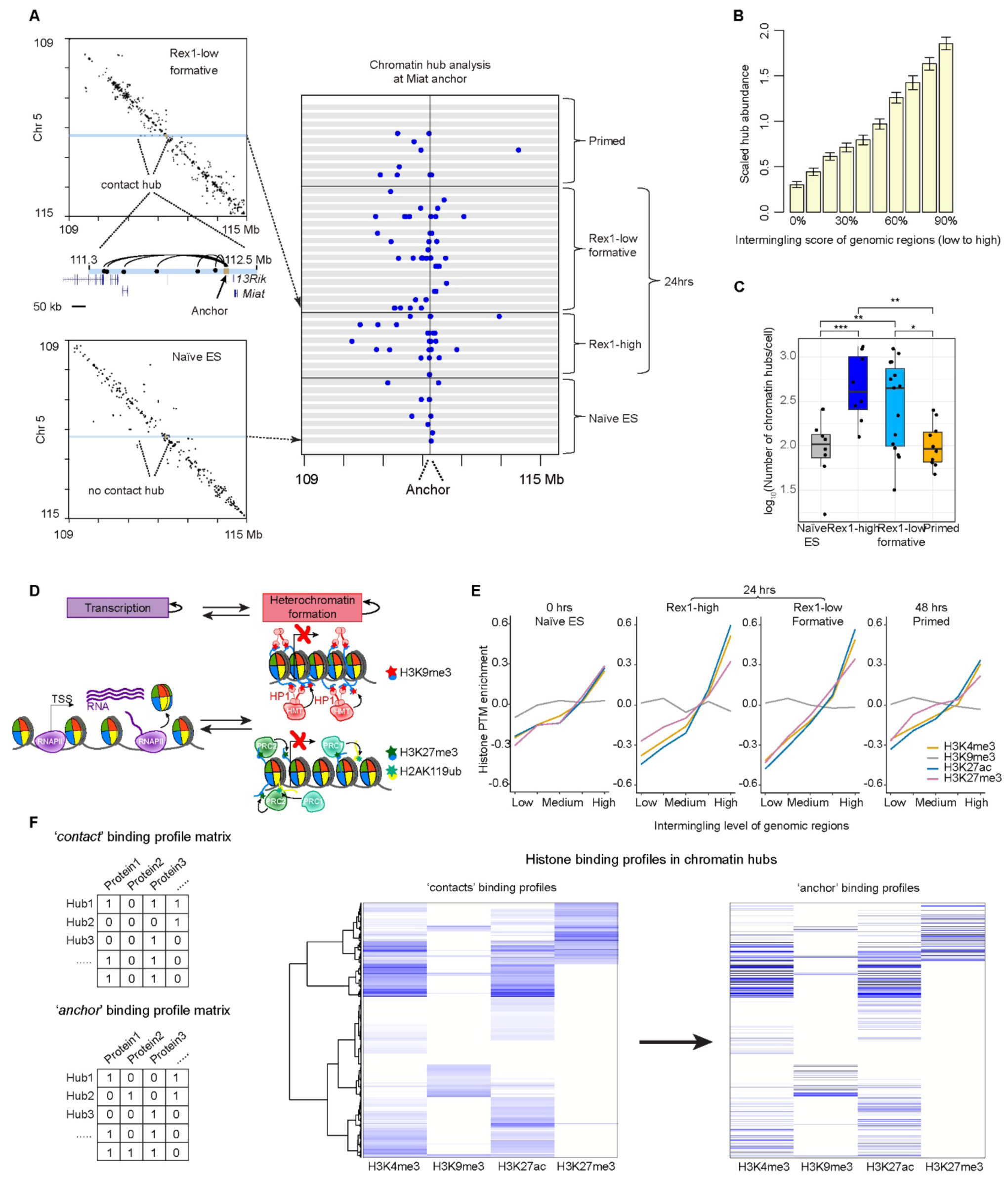
Multiway contacts in 3D genome structures of single cells identify chromatin hubs. **A**) (Left) an example of a multiway chromatin hub which mainly forms in 24 hr cells whose ‘anchor’ is nearby the *Miat* ncRNA gene. (Right) Contacts made from the anchor region near the *Miat* ncRNA locus in different cells and stages of differentiation – each grey horizontal box shows the same small section of the contact map (see Left) from a different individual cell. **B)** After dividing the genome into 10 equal-sized groups according to the level of chromosome intermingling, the abundance of hubs was found for each group and scaled by the average number of hubs across all the groups. (The error bars show a bootstrap estimate of the standard deviation of hub abundance within each group obtained from 2000 random samples of 30% of the data.) **C)** Boxplots showing the number of multiway chromatin hubs identified in individual single cells at different stages of differentiation. The midline is the median value, the lower and upper edges of the box represent the 1st and the 3rd quartiles, and the whiskers represent the minimum and maximum values. FDR adjusted Mann Whitney U test *p*-values are also shown: * - *p* < 0.05; ** - *p* < 0.01; *** - *p* < 0.001. **D)** Transcription and heterochromatin formation are self-reinforcing processes leading to the deposition/modification of nucleosomes with either active (H3K4me3, H3K27ac) or repressive (H3K9me3/H3K27me3) histone marks and heritably expressed or silenced states. **E)** H3K4me3, H3K27ac and H3K27me3, but not H3K9me3, modified nucleosomes are found in highly intermingled regions of the A compartment in formative state cells. In this analysis the A compartment of the genome was divided into 5 groups according to the intermingling level. The relative abundance of each type of histone modification was then compared to the average levels, and a Fisher’s Exact test was calculated when comparing the density in the highest and the lowest intermingled regions (*p* < 10^-15^ for all the modifications except H3K9me3). **F)** (Left) Schematic illustrating the approach used to analyse ‘anchor’ or ‘average contact’ histone protein binding profiles in individual hubs in single cells. [NB – In practice the actual number of ChIP-seq reads was used as opposed to a binary (0, 1) distinction of histone binding or not binding.] (Middle) An average protein binding profile matrix was calculated for all the contacts in a hub, which were then ordered using hierarchal clustering. (Right) By keeping the row order the same each line continues to represent the same chromatin hub, but now shows the histone protein binding profile at the anchor in that hub (as opposed to the contacts).

Interestingly, despite the enormous variation in 3D genome structure from cell-to-cell, and the way chromosomes intermingle with each other, we found that some loci form chromatin hubs in multiple cells (including the example in Figure 3A). This finding that the same anchor locus can make different contacts in different cells, suggests that the 3D structures may provide snapshots of a dynamic process(es) that leads to the generation of these chromatin hubs. Further analysis showed that inter-chromosomal intermingling and the formation of chromatin hubs are very significantly associated with each other (Figure 3B). As with inter-chromosomal intermingling, in both Rex1-high and low cells there is a very significant increase in the abundance of multiway chromatin hubs in 24 hr cells compared to either the naïve or primed states (Figure 3C). Indeed, multiway chromatin hubs are largely a feature of the 24 hr state and in both Rex1-high and low cells they mostly involve contacts between two loci in the A compartment (A-A contacts) or between loci in the A and B compartments (A-B contacts) (Supplementary Figure 2). Strikingly, inspection of the 3D structures showed that multiway chromatin hubs tend to form at the edge of the intermingled regions – i.e. at the interface between the region of the chromosome that is buried and the regions that are intermingled with other chromosomes (Figure 2C). Whilst the multiway chromatin hubs that are formed and the chromosomes that intermingle with each other varies from cell-to-cell, the regions of a chromosome that intermingle are much more consistent – with the average intermingling scores from as few as three or five cells clearly matching the population average (Supplementary Figures 3A,B). Multiway chromatin hubs fall into two groups whose maximum sequence span of intra-chromosomal contacts is either less than ∼1 Mb or many Mb in length (Supplementary Figure 3C). We hypothesised that while the shorter range contacts may form through DNA loop extrusion^28,34,35^, the much longer distance contacts may be generated through transcription or the formation of Polycomb bodies^36-38^.

Thus, 3D genome structure undergoes a very significant transition in the 24 hr state (in both Rex1-high and low cells), with a substantial increase in inter-chromosomal intermingling and the formation of multiway chromatin hubs, which are typically found at the boundary between the buried and intermingled regions of a chromosome.

### Regions of inter-chromosomal intermingling and multiway chromatin hubs are enriched in markers of active gene expression and Polycomb-mediated heterochromatin

Transcription and heterochromatin formation are opposing processes, whose balance at a particular gene results in the deposition or modification of nucleosomes with either active (H3K4me3, H3K27ac) or repressive (H3K9me3, H3K27me3) histone marks, and heritable states of either active gene expression or silencing, respectively (Figure 3D). The changes in A/B compartmentalisation at 24 hrs (Figures 1C,E) suggested that the alterations in 3D genome structure might involve a reorganisation of the interactions between eu- and hetero-chromatic regions of the genome. This prompted us to carry out ChIP-seq experiments in naïve ES, 24 hr and primed cells to study the binding of post-translationally modified histones to identify different chromatin states. We then mapped the resulting data onto the single nucleus 3D genome structures and found that marks of active enhancers and promoters (H3K27ac, H3K4me3) and Polycomb-mediated heterochromatin (H3K27me3) were all highly associated with inter-chromosomal intermingling in the 24 hr state, but this was much less noticeable in naïve or primed cells (Figure 3E).

Hierarchical clustering of histone binding profiles in individual chromatin hubs in single cells showed that ‘*anchors*’ and ‘*contacts*’ with very similar chromatin states cluster together in the same hub: some hubs were found to bear marks of active transcription (H3K4me3, H3K27ac), some had marks of Polycomb-mediated heterochromatin (H3K27me3), and some had marks of both (Figure 3F). The smaller number of hubs bearing the H3K9me3 mark were not enriched above background: in addition, the association of H3K27me3, but not H3K9me3, with inter-chromosomal intermingling in the 24 hr state (Figure 3E) prompted us to focus on changes in the formation of Polycomb-mediated heterochromatin.

We found that in the 24 hr state there is a global reduction in H3K27me3 levels – this decreases throughout the promoter and gene body and then builds up again at the promoter in primed cells (Figure 4A), suggesting a loss of Polycomb-mediated heterochromatin and its subsequent reestablishment^39^. We classified genes according to how H3K27 methylation levels change at their promoters and identified four major groups, whose Gene Ontology (GO) terms were all found to be significantly related to embryonic development, cell differentiation and cell-cell signalling. Interestingly, the top hits for Group 1 genes, which follow the global changes in H3K27me3 levels (see Figure 4A), all relate to neuroectoderm development – e.g. the regulation of neural precursor cell proliferation, sensory system development and synapse organisation (Figure 4B). In Group 2 genes, promoter H3K27me3 levels increase throughout the time course despite the global decrease in the 24 hr state (Figure 4A). This group can be split into two: group 2A consists of genes whose promoter H3K27me3 levels increase in both the 24 hr and primed states, whilst in Group 2B the levels increase more modestly in the 24 hr state (than in Group 2A) and then remain stable in the primed state. We found that GO terms for Group 2 genes do not relate to neuroectoderm development, but are instead biased towards the meso/endodermal lineages – e.g. embryonic organ/skeletal system development (Group2A) and tissue remodelling (Group2B) (Figure 4B). Finally, GO terms for Group 3 genes are most significantly related to the organisation of intermediate filament-based processes and fibre organisation – processes that could be of importance in early cell differentiation. It is also worth noting that GO terms for Groups 1 and 2A genes relate to distinct signalling pathways (MAPK and BMP, respectively) (Figure 4B). (A complete list of the genes in the different H3K27me3 groups can be found in Supplementary Table 4, and the GO terms relating to these different groups of genes can be found in Supplementary Tables 5-8.)

**Figure 4.**
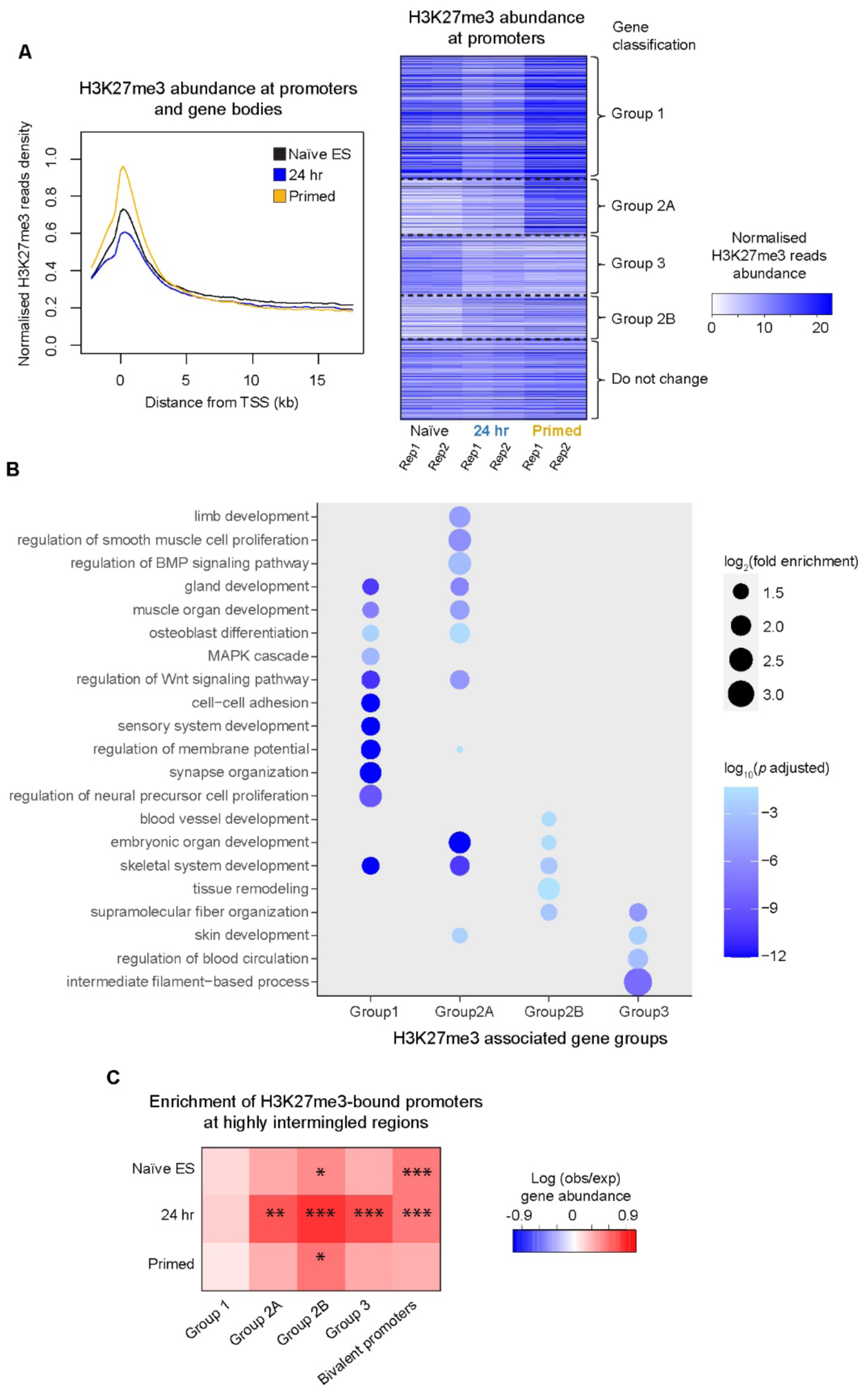
There is a global loss of H3K27me3 in 24 hour state cells. **A)** (Left) Pileup analysis of histone H3K27me3 binding to genes, plotted in 200 bp bins from 2 kb upstream of the transcription start site (TSS) across the gene body. (Right) When looking at promoters bound by nucleosomes with this histone modification, genes can be hierarchically clustered into four different groups depending on how their H3K27me3 binding levels change during the time course. **B)** Gene Ontology (GO) enrichment analysis for the different groups of H3K27 methylated genes – the dot-size scales with Log (fold enrichment), and FDR adjusted *p*-values (Fisher’s exact test) are colour coded in different shades of blue. **C)** The enrichment of H3K27me3-bound promoters within genomic regions having high levels of A compartment inter- chromosomal intermingling.

Both Group 2 and Group 3 promoters – which in 24 hr cells have increasing and decreasing levels of H3K27me3, respectively – are enriched in regions of high inter-chromosomal intermingling (Figure 4C). We identified bivalent promoters (marked by both H3K4me3 and H3K27me3) and tracked their formation from, and resolution into, monovalent promoters (either H3K4 or H3K27 methylated) during the time course (Supplementary Figure 4A). Bivalent promoters were also found to be particularly enriched in regions of high inter-chromosomal intermingling in naïve ES and 24 hr cells (Figure 4C). Furthermore, promoters that are bivalent at all three stages in the time course were found to be enriched in Group 1 genes, whilst those which form in the 24 hr state and are maintained in the primed state are enriched in Group 2B genes (Supplementary Figure 4B). Finally, bivalent promoters that form only in naïve ES or primed cells are enriched in either Group 3 or Group 2A genes, respectively (Supplementary Figure 4B).

Thus, the 24 hr state is characterised by an alteration in H3K27me3 levels at promoters that are found in regions of high inter-chromosomal intermingling and chromatin hubs, suggesting a switch in the balance between transcription and the formation of Polycomb-mediated heterochromatin. Moreover, genes whose H3K27me3 levels change in different ways are associated with alternative developmental lineages – suggesting that changes in gene expression/Polycomb-mediated heterochromatin formation might prime them for differential expression later in differentiation.

### Regions of inter-chromosomal intermingling and multiway chromatin hubs are enriched in genes that are required for the exit from naïve pluripotency

To explore changes in gene expression we carried out single cell RNA-seq experiments in naïve ES, 24 hr and primed cells (Figures 5A,B). We found that there is a good correlation between the classification of groups of genes based on changing levels of H3K27 methylation and the levels of expression in the scRNA-seq data. In particular Group 2A and Group 3 genes, which respectively have increasing and decreasing levels of H3K27me3 at their promoters, are significantly enriched amongst genes that are down- and up-regulated during the time course (Supplementary Figure 4C). (A complete list of the genes in the different scRNA-seq groups can be found in Supplementary Table 9.) A small proportion of genes (1.8%) whose expression levels increase in the 24 hr state move from the B to the A compartment upon differentiation, but most (74.4%) are constitutive A compartment genes. Interestingly, we found that genes that are either up-, down- or transiently upregulated in the 24 hr state are significantly enriched in regions of high inter-chromosome intermingling in the A compartment (Figure 5C). Moreover, consistent with the 3D structures which show that the formation of multiway chromatin hubs and inter-chromosomal intermingling is very closely related (Figures 2C & 3B), differentially expressed genes are also significantly enriched in multiway chromatin hubs (Figure 5C).

**Figure 5.**
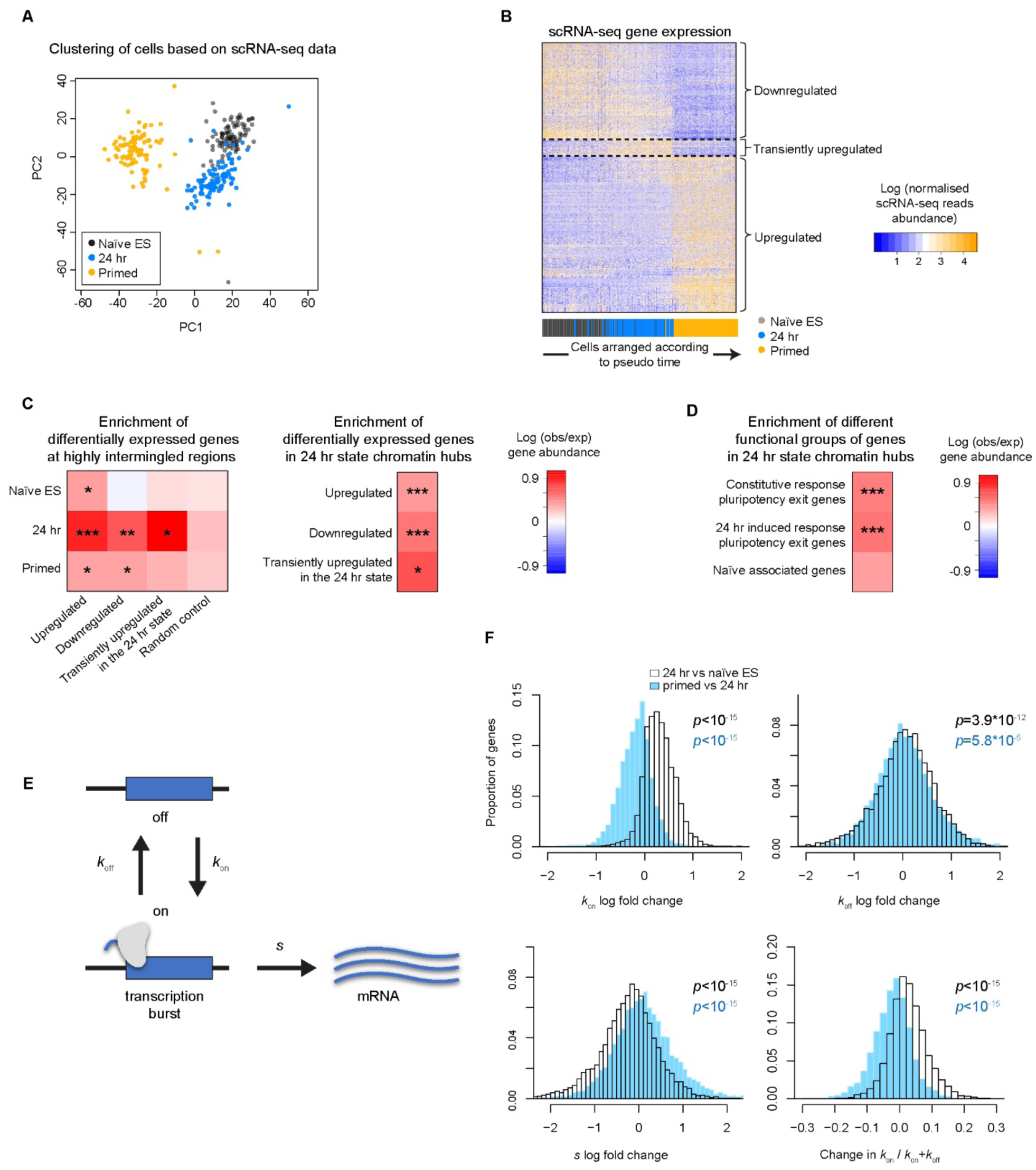
There is a global increase in transcriptional bursting in 24 hour state cells. **A)** After PCA analysis single cell (sc) RNA-seq data (0, 24 and 48 hrs after induction of differentiation) can be separated using k-means clustering. **B)** Expression profile for differentially expressed genes in the scRNA-seq data (each cell is represented by a different column). The cells were ordered according to pseudo time (see Methods), with the naïve ES, 24 hr and primed cell datasets being coloured black, blue and yellow, respectively in the bar chart. Differentially expressed genes (individual rows) were hierarchically clustered to identify three categories of genes whose expression levels change during the time course. **C)** The enrichment of differentially expressed genes at different time points within either (Left) genomic regions having high levels of A compartment inter-chromosomal intermingling or (Right) chromatin hubs. **D)** The enrichment of pluripotency exit and naïve-associated genes in 24 hr chromatin hubs. [Very similar results were obtained when analysing the separate Rex1-high and -low (formative state) cells (data not shown).] The colours shown in the heatmaps in C) and D) indicate the log fold change in gene abundance – we compared the top 30% most intermingled A compartment regions with the bottom 30%, and contacts observed within 24 hr state chromatin hubs to the genome wide distribution (see Methods). FDR adjusted empirical *p*-values are also shown (* - *p* < 0.05; ** - *p* < 0.01; *** - *p* < 0.001). **E)** The model used to estimate kinetic parameters from the scRNA-seq data^42^. 1/*k*off and 1/*k*on describe the average time a gene spends in the active and inactive states, respectively, whilst *s* is the transcription rate. **F)** Histograms showing the log fold changes in *k*on, *k*off and *s*, as well as changes in the average fraction of time that genes spend in the active state (*k*on/(*k*on+*k*off) during the time course. (Wilcoxon paired tests *p* < 10^-15^ for all *k*on, *s,* and *k*on/(*k*on+*k*off) comparisons of naïve ES vs 24 hr, and 24 hr vs primed cells. The values for *k*off were *p* < 3.9*10^-12^ when comparing naïve ES and 24 hr cells, and 5.8*10^-5^ when comparing 24 hr and primed cells.)

In previous work we carried out a comprehensive mutagenic screen to identify regulators that are important for the exit from naïve pluripotency. We then used transcriptomic information from 73 differentiation defective knockout ES cell lines to identify gene-sets that are functionally relevant for the exit from naïve pluripotency^40^. These gene-sets fall into two groups: *1)* genes which are deregulated by a KO in both 2i/LIF and 24 hr conditions – termed ‘*constitutive response*’ genes and; *2)* genes which are not (or only weakly) deregulated by a KO in 2i/LIF conditions but are significantly deregulated in the 24 hr state – termed ‘*24 hr-induced response*’ genes. We also identified an extended group of genes that are closely associated with the naïve ES cell state (‘*naïve associated*’ genes), which we found were very enriched in Group 2A genes. All three groups of functionally important genes were found to be enriched in 24 hr chromatin hubs – the two groups of pluripotency exit genes very significantly so (Figure 5D).

Thus, genes which are of potential functional relevance for the exit from naïve pluripotency are enriched in chromatin hubs/regions of high inter-chromosome intermingling in the A compartment in 24 hr state cells.

### There is a global change in transcriptional kinetics and a reorganisation of enhancer-promoter interactions in the 24 hr transition state

Previous work has shown that a loss of PRC1-mediated gene silencing leads to an increase in the activation rate *kon* (or burst frequency) of promoters ^41^. We employed a minimal model of transcriptional bursting^42^ to study the expression kinetics of individual genes using the scRNA-seq data (Figure 5E). Surprisingly, when looking at all active genes – which are concentrated in the A compartment – we found that there is a very significant global increase in the burst frequency *kon* in the 24 hr state compared to naïve ES and primed cells (Figure 5F). In addition to genes turning on more frequently the rate of transcription *s*, i.e. the amount of RNA made each time a gene switches on, decreases (Figure 5F). We found that genes in Groups 1, 3 and 2B, whose H3K27me3 levels either decrease or only increase slightly at 24 hrs, follow this global trend of increased transcriptional bursting (Supplementary Figure 5). By contrast, transcriptional bursting does not increase in Group 2A genes, whose H3K27me3 levels go up across the time course (Supplementary Figure 5).

We hypothesized that in the 24 hr transition state the disruption of Polycomb-mediated heterochromatin/genome structure, together with the increase in transcriptional bursting, might allow new enhancer-promoter interactions to form through a ‘transcriptional stirring’ mechanism^43^. After assigning the activity of individual enhancers and promoters in naïve ES, 24 hr state and primed cells using the ChIP-seq data, we therefore analysed changes in enhancer-promoter interactions. We found that there is a significant enrichment of interactions between down-regulated genes and weakening enhancers in 24 hr state multiway chromatin hubs (Figure 6A). There is also, however, a much more significant enrichment of interactions between up-regulated genes and emerging enhancers (Figure 6A). This was most noticeable for interactions between promoters in hub anchors and enhancers in hub contacts. Furthermore, in 24 hr state multiway chromatin hubs we found that promoters of naïve associated genes tend to contact weakening enhancers in cells that have progressed into the formative Rex1-low state (Figure 6B). In contrast, both the constitutive and 24 hr-induced response genes required for pluripotency exit tend to contact emerging enhancers in Rex1-low cells (Figure 6B). These trends were much less obvious in the Rex1-high state, where cells are still exiting naïve pluripotency.

**Figure 6.**
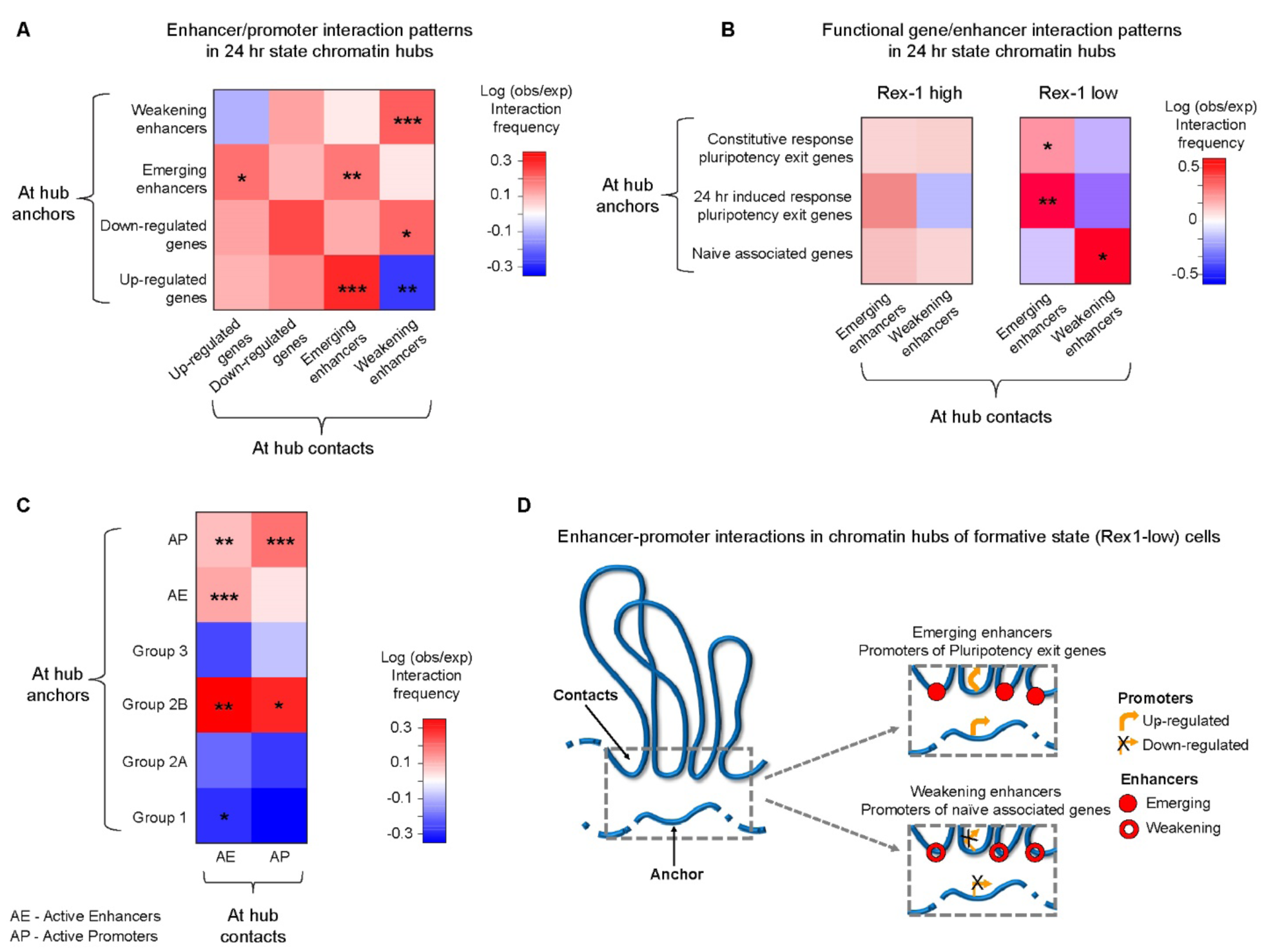
Enhancer-promoter interaction patterns in formative state chromatin hubs. **A)**, **B)** and **C)** Heatmaps showing the observed vs expected interaction frequency of contacts that form within 24 hr state chromatin hubs between either up-/down-regulated promoters and emerging/weakening enhancers (A), or between pluripotency exit/naïve associated promoters and emerging/weakening enhancers (B), or between the different groups of H3K27 methylated genes and active enhancers/promoters (C). In A), B) and C) the colours shown in the heatmaps indicate the log fold change in interaction frequency compared to the randomly expected distribution of the number of hub contacts between regulatory elements. FDR adjusted empirical *p*-values are also shown (* - *p* < 0.05; ** - *p* < 0.01; *** - *p* < 0.001). **D)** Schematic illustrating the finding that upregulated genes contact emerging enhancers and, conversely, down-regulated genes contact weakening enhancers in long-range multiway chromatin hubs.

Because multiway chromatin hubs tend to be at the interface between the buried core and the regions of the chromosome that are intermingled with other chromosomes (i.e. not on the surface, see Figure 2C) we find relatively few *inter-chromosomal* interactions in hubs. Thus, the enrichments of interactions between genes and enhancers in 24 hr state multiway chromatin hubs are mostly *intra-chromosomal*. However, given the close association between multiway chromatin hubs and inter-chromosomal intermingling (Figures 2C & 3B), we wondered whether the latter might also lead to a reconfiguration of *inter-chromosomal* enhancer-promoter interactions. Indeed, we find that both constitutive and 24 hr-induced response genes required for pluripotency exit also tend to contact emerging *inter-chromosomal* enhancers in formative state (Rex1-low) cells (Supplementary Figure 6), although the enrichments are weaker than those observed in multiway chromatin hubs. As with multiway chromatin hubs, this trend wasn’t observed in the less differentiated Rex1-high state.

Taken together, these results show that Rex1-low cells may have progressed into the ‘formative’ state because genes required for differentiation have established long-range interactions with emerging enhancers whereas, conversely, naïve associated genes are interacting with weakening enhancers. This switch in the pattern of enhancer-promoter interactions is not properly established in Rex-l high cells, perhaps explaining why Rex1- high (but not Rex1-low) cells can revert to the pluripotent state. We also found that there is a very significant enrichment in the interactions between promoters of the Group 2B mesodermal/endodermal genes, e.g. *Edar*, *Fstl3* and genes in the Hox clusters, with active enhancers and promoters in 24 hr state chromatin hubs (Figure 6C). This further suggests that structural changes in the 24 hr state may set up Group 2B enhancer-promoter interactions, priming them for future lineage-specific expression.

### Formation of the formative state structure is controlled by DNMT3a/3b and TET1

How might the structural changes we observe in the 24 hr state be controlled? In naïve ES cells levels of CpG methylation are low, and during differentiation they are regulated by the opposing activities of the de novo DNA methyltransferases DNMT3a/3b and the 5-methyl-cytosine dioxygenase TET1^44,45^ (Figure 7A). Previous work suggested that an increase in DNA methylation by DNMT3a/3b could lead to reduced PRC1 recruitment^46-48^ and lower levels of H3K27me3^38^, consistent with the decrease of H3K27 methylated promoters in 24 hr state cells (Figure 4A). Significantly, in-nucleus Hi-C experiments of *Dnmt3a/3b-*dko mESCs^49,50^ showed that there was very little change in either contact probability (Figure 7B) or A/B compartmentalisation 24 hrs after exit from the naïve state (Figure 7C) – i.e. formation of the 24 hr state genome structure appears to be prevented in the absence of de novo DNA methylation.

**Figure 7.**
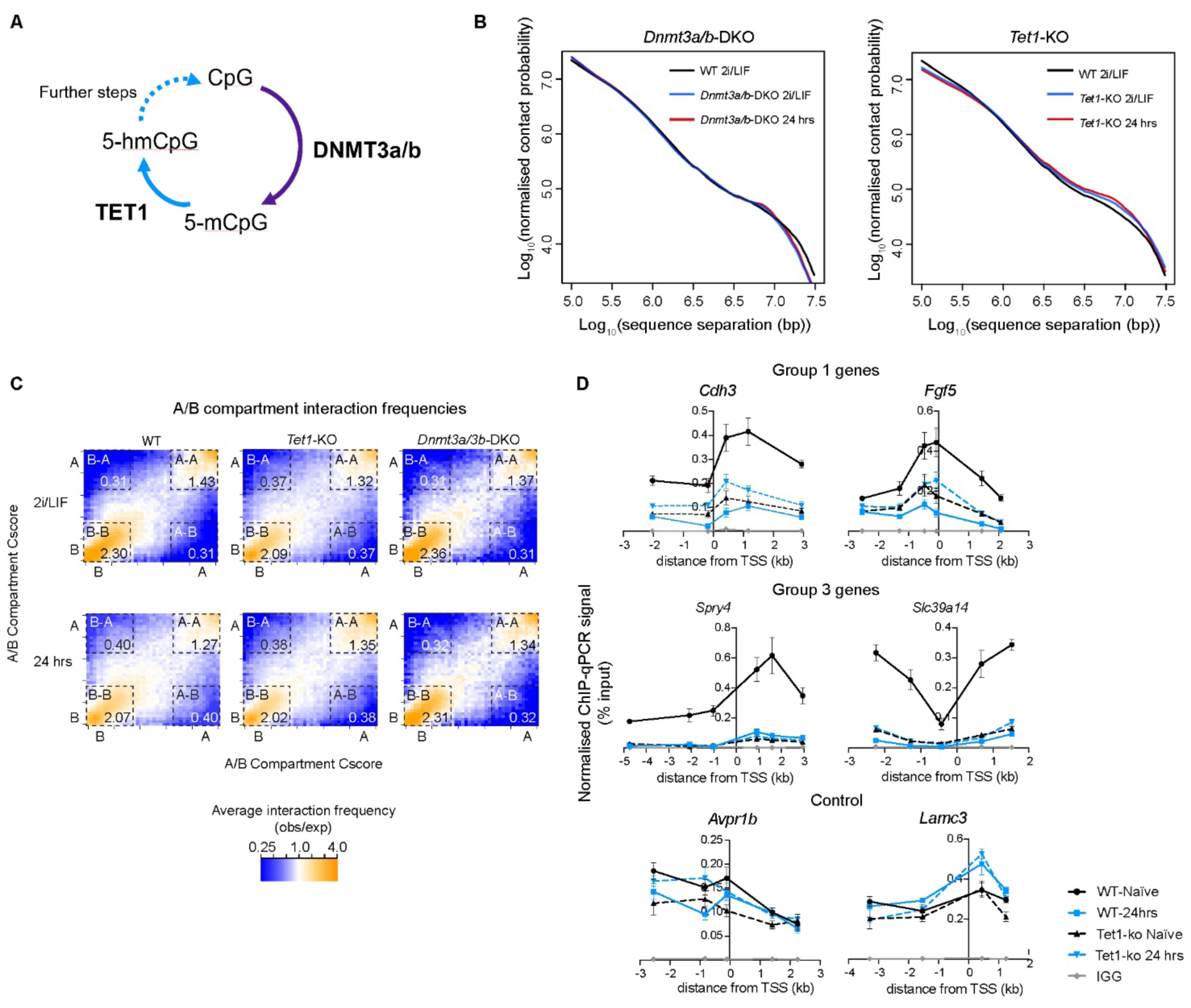
Progression to the formative state genome structure is controlled by DNMT3a/3b and TET1. **A)** Scheme showing how DNMT3a/b and TET1 respectively methylate and demethylate CpG sequences to control the levels of DNA methylation at CpG islands. **B)** Plots of log10 contact probability against log10 intra- chromosomal sequence separation determined using in-nucleus Hi-C experiments show that: (left) the double knock out of *Dnmt3a/3b* prevents transition to a formative state nuclear structure; whilst (right) the formative state structure is generated in the *Tet1* knock out. **C)** Saddle plots showing the normalised average A/B compartment interaction frequencies between pairs of 25 kb bins, arranged by their A or B compartment Cscore’s (see Methods). The numbers in the corners show the average normalised Cscore within the boxed A/B compartment regions. **D)** Comparison of H2K27me3 levels in the *Tet1* knock out and its background wild-type cell line^40,54^ across the promoters of two representative Group 1 and 3 genes as determined using ChIP-qPCR experiments in naïve ES and 24 hr cells. The data are plotted as the mean of three biological replicates and the error bars represent standard error. The controls are genes whose H3K27me3 levels did not change significantly in the ChIP-seq experiments across the time course.

The TET1 enzyme binds to unmethylated CpGs in naïve mESCs and co-localises with most PRC1/2 target genes^44,45^. Strikingly, consistent with previous work showing that *Tet1* deletion reduces the deposition of both H2AK119ub and H3K27me3^51^, we found that in 2i/LIF conditions genome structure in the *Tet1*-ko^40^ closely resembles that of wild-type cells in the 24 hr state – as evidenced by similar Hi-C contact probabilities and weakened A-B compartment insulation (Figures 7B,C). This suggested that *Tet1* deletion leads to reduced Polycomb-mediated heterochromatin formation in naïve ES cells. We confirmed this by showing that in 2i/LIF conditions H3K27me3 levels at the promoters of Group 1 and 3 genes in the *Tet1*-ko are reduced to levels similar to that seen in the wild-type background at 24 hrs (Figure 7D). Moreover, allowing the cells to differentiate for 24 hrs caused little further change in either contact probability or A/B compartmentalisation showing that the structural changes had already occurred (Figures 7B,C). Importantly, this experiment suggested that the structural changes we observe in the formative state are not due to changes associated with differentiation – because in the *Tet1*-ko the changes are observed in 2i/LIF conditions which maintains cells in a pluripotent state^52^.

## Discussion

The 3D genome structures show that as naïve ES cells become specified towards different cell fates they go through a profound structural transformation in the formative state, with the formation of active (H3K4me3/H3K27ac), repressed (H3K27me3) and bivalent (H3K4me3/H3K27me3) long-range multiway chromatin hubs and very significant inter-chromosomal intermingling. We find that genes whose expression levels are changing become located in these long-range multiway chromatin hubs with, most notably, genes required for pluripotency exit establishing *intra-chromosomal* interactions with emerging enhancers (see Figure 6D). There is also a significant, albeit less marked, increase in interactions between pluripotency exit gene promoters and emerging *inter-chromosomal* enhancers as a result of the increased intermingling between different chromosomes in the formative state. Furthermore, we found that in 24 hr cells promoters of the Group 2B meso/endodermal genes establish contacts with active enhancers and promoters in multiway chromatin hubs long before they are expressed, suggesting that the formative state structure may also prime genes for activation later in development.

In previous studies increased DNA methylation has been found to reduce PRC1 recruitment^46-48^ leading to lower levels of H3K27me3^38^. A global drop in histone H3K27me3 levels in the formative state suggested that the increased chromosomal intermingling and reconfiguration of enhancer-promoter contacts could result from decreased interactions between regions of the genome bound by the Polycomb complexes PRC1/2. The changes in Hi-C contact probability and A/B compartmentalisation in the *Dnmt3a/3b* and *Tet1* knockouts, together with the observation of decreased H3K27me3 levels in naïve cells in the *Tet1*-ko, strongly suggest that the changes in 3D structure that we observe in formative state cells are driven by increased *de-novo* DNA methylation. Consistent with previous studies showing that a loss of PRC1 binding leads to increased transcriptional bursting (*i.e.* increased *k*on)^41^ we found that there is also a global increase in *k*on in the formative state.

It is not clear how the formation of multiway chromatin hubs and inter-chromosomal intermingling are mechanistically linked. At the outset of this work we wondered whether the structural changes we observe could result from the altered activity of a particular chromatin regulator that leads to global changes in chromatin dynamics – this in turn could facilitate the reconfiguration of enhancer-promoter interactions. We also wondered whether globally altered chromatin dynamics might explain the previously observed differences in the material properties of 24 hr nuclei^8^. However, our results suggest that there is no global change in either eu- or hetero-chromatin dynamics during this time course of differentiation (data not shown). We therefore favour the idea that the loss of PRC1 binding leads to locally increased transcriptional bursting, which in turn locally increases enhancer/promoter dynamics through a transcriptional stirring mechanism^43^. This locally increased chromatin dynamics could directly lead to the increased inter-chromosomal intermingling in the 3D structures, which would be consistent with it being mainly observed in the A (euchromatic) compartment (Figure 2A). It might also enable the formation of multiway chromatin hubs through facilitating the establishment of new interactions between regions of the genome having similar chromatin states – in this regard the similar chromatin state of anchors and contacts in hubs (Figure 3F) is noteworthy.

Our finding that the formation of long-range multiway chromatin hubs and increased intermingling in the formative state leads to a reconfiguration of enhancer-promoter interactions as the genome goes through a dramatic transient structural reorganisation suggests a hitherto under appreciated role for much longer-range enhancer-promoter interactions. Up to now most studies of enhancer-promoter contacts have focussed on relatively short-range genomic interactions – e.g. between genomic sequences that are either within the same TAD (∼1 Mb) or even between a promoter and its nearest active enhancer in the linear chromosome sequence. This makes sense because sequences that are tethered nearby to each other are much more likely to interact, and Cohesin will transiently bring sequences together that are normally further apart as DNA loops are extruded (leading to the observed increase in contact probability within TADs). However, by showing that interactions between emerging enhancers and pluripotency exit genes have become established in cells that have progressed into the formative state (Rex1-low), but not in cells that are slightly less differentiated (Rex1- high), we provide evidence that the formation of unusually long-range enhancer-promoter interactions in multiway chromatin hubs and in the regions of chromosomes that intermingle with each other may also be functionally important. Our results lead to a model whereby those cells whose promoters first generate suitable enhancer interactions (as a result of the initial transcriptional bursting) become the first to attain the formative Rex1-low state. It is not clear whether direct contacts are required for enhancers to activate promoters, or whether activation occurs through a diffusible signal, or whether the clustering of enhancers and promoters in phase-separated hubs leads to gene activation^11^, but we note that our results would be compatible with any of these mechanisms. It is also not clear whether Cohesin-meditated interactions could play a role in the formation of the very long-range interactions that we see in the formative state. Our preliminary analysis suggests that CTCF/Cohesin are often associated with the shorter-, but not the longer-range contacts in multiway chromatin hubs (data not shown), arguing that processes other than Cohesin-mediated loop extrusion lead to their formation. In this regard the large number of active multiway chromatin hubs (marked by H3K4me3 and H3K27ac, see Figure 3F) is noteworthy, and it suggests that interactions between actively transcribed regions may be an important factor. However, Cohesin could bind to stabilise hubs as shown previously for TAD-like domains in the parental to embryo switch^53^.

We find that chromosomes intermingle with different partners and that different multiway chromatin hubs are established in different individual cells – i.e. the enhancers that interact with a particular promoter varies from cell-to-cell. At first sight this result may seem surprising, but we think it could be important in the context of the limited time available for chromosomes to unfold in the G1 phase of the cell cycle following mitosis. In that limited time chromosomes cannot unfold and form a consistent 3D structure, and a promoter cannot extensively search the nucleus to find a particular enhancer. Rather, it is much more likely that genes will be stably expressed if a promoter can be activated through the interaction with different compatible enhancers in different cells. The detection of long-range contacts where promoters establish interactions with different enhancers in different single cells isn’t possible when using 3C based experiments carried out on populations of cells. Up to now we have therefore lacked the ability to detect such interactions, but single cell Hi-C and 3D structure determination provides the means to study long-range enhancer-promoter interactions genome-wide. The 3D genome structures of mouse ES cells in the naïve, formative and primed states of pluripotency show that very rapid changes in chromatin structure and enhancer-promoter interactions (over 1-2 cell cycles) can be linked to altered gene expression, and they raise the intriguing possibility that other changes in either cell fate or cell state could involve a similar structural reorganisation.

## Acknowledgements

We thank Tessa Kretschmann for preparing the figures for publication, Grace Ding for carrying out preliminary snHi-C experiments of Rex1-low (formative state) cells, Austin Smith for the *Dnmt3a/b* double knockout cell line and the CSCI flow cytometry (Andy Riddell) and DNA sequencing (Maike Paramor and Vicki Murray) core facilities. This research was funded by the EU FP7 Integrated Project “4DCellFate” (277899), the Medical Research Council (MR/P019471/1) and the Wellcome Trust (206291/Z/17/Z). We also thank the MRC (MR/R009759/1 to BDH, and MR/M010082/1 to EDL) and the Isaac Newton Trust (17.24(aa) to BDH) for project grant funding, as well as the Wellcome Trust/MRC for core funding (203151/Z/16/Z) to the Cambridge Stem Cell Institute.

## Author contributions

D.L ., X.M., and Y.C. carried out the Hi-C experiments. T.J.S. with assistance from W.B. developed the software for single cell Hi-C processing and genome structure calculation and visualisation. N.R. and B.D.H. carried out the ChIP-seq and ChIP-qPCR experiments while B.M. and B.D.H. carried out the single-cell RNA-seq experiments. X.M. and W.B. processed the raw ChIP sequencing data and R.R. processed the raw RNA sequencing data. M.L. created the haploid *Rex1*-dGFP reporter ES cells, and with A.L. created the *Tet1*-ko ES cells and carried out preliminary H3K27me3 ChIP-seq experiments. A.J. carried out differentiation experiments and studies of chromatin dynamics. X.M. and D.H. carried out computational analysis. D.L., X.M. and E.D.L. designed the experiments, analysed the results and wrote the manuscript with contributions from all the other authors.

## Data availability

The Hi-C, ChIP-seq and single cell RNA-seq datasets reported in this study are available from the Gene Expression Omnibus (GEO) repository under accession code GSE214264.

## Biological materials

All constructs and cell lines are available upon request.

## Code availability

All the code used is freely available as indicated in the Methods.

## Methods

### Cell lines, cell culture and differentiation protocol

The mouse ES cell lines employed have all been described previously (see Table below). The haploid mouse embryonic stem (mES) cell line containing the Rex1-GFPd2-IRES-BSD reporter was sorted every 4-6 passages to enrich for haploid cells^55^.

| Cell line | Genotype | Reference |
| --- | --- | --- |
| Haploid | Rex1GFPd2-IRES-BSD | Leeb *et al.*, *Cell Stem Cell*, 2014, **14**(3): 385–393 |
| Wild-type (Diploid) | RC9 | Li *et al.*, *Cell Rep.*, 2018, **24**(2): 489–502 |
| *Tet1*-ko | *Tet1*-ko/RC9 | Lackner *et al.*, *EMBO J*, 2021, **40**(8): e105776 |
| *Dnmt3a/b*-dko | *Dnmt3a/b*-dko/RC9 | Li *et al.*, *eLife*, 2017, **6**: e23468 |

For routine cell culture, cell lines were maintained in N2B27 media supplemented with 2i inhibitors and mouse leukemia inhibitory factor (mLIF)^52^. 50 % DMEM/F-12 medium (Gibco, 21041025) and 50 % Neurobasal™ Medium (Gibco, 12348017) was supplemented with 1 x N2 to a final concentration of 2.5 µg/ml insulin (provided in-house by the Cambridge Stem Cell Institute), 0.5x B-27™ Supplement (Gibco, 17504044), 1x minimum essential medium non-essential amino acids supplement (Sigma-Aldrich, M7145), 2 mM L-glutamine (Life tech, 25030024), 0.1 mM 2-mercaptoethanol (Life tech, 21985023), 2i inhibitors (1 µM PD0325901, 3 µM CHIR99021) and 10 ng/ml mLIF (provided by the Biochemistry Department, University of Cambridge). Cells were routinely screened for mycoplasma contamination.

For differentiation experiments we prepared samples as previously described^5^. Cells were plated at a density of 10,000 cells/cm^2^ in N2B27 with 2i and no LIF to avoid delays in the onset of differentiation^56^. After 24 hrs the two inhibitors were removed and cells were grown in N2B27 only media for a further 24 or 48 hrs to generate Rex1-high/Rex1-low (formative) or primed state cells. To obtain pure Rex1-high and -low fractions, cells at the 24 hr stage were FACS sorted with gates set to collect those with the 25% highest and lowest GFP expression. Naïve ES state cells were plated at the same density and grown in N2B27 with 2i and LIF for 48 hrs.

### Population Hi-C protocol and samples

Population Hi-C processing to generate ligated contacts was carried out as follows. Four-five million cells were fixed for 10 min in 14 ml of 2% formaldehyde in growth media at room temperature and then the reaction was quenched by the addition of 1 ml of 2 M glycine. Nuclei were then extracted by incubating cells on ice for 30 min (inverting every 10 min) in 10 mM Tris pH 8.0, 10 mM NaCl, 0.2% NP-40 (IGEPAL CA- 630) and protease inhibitor cocktail (Roche). After transferring nuclei to a 1.5 ml Eppendorf tube, the pellet was resuspended in 50 ul 0.5% SDS before incubation for 5 min at 62°C. After incubation, 145 ul of Milli-Q water followed by 25 ul of 10% Triton-X100 were added and the nuclei were incubated again for 15 min at 37°C with gentle agitation. The DNA was digested overnight at 37°C with gentle agitation after adding 25 µl of 10x NEBuffer3 and 5 ul of 25 U/ul MboI (NEB). To biotin end-fill the DNA a 50 ul master mix containing the following was added: 0.3 mM each of dCTP, dGTP, dTTP and biotin-14-dATP (Jena Biosciences) together with 40 U DNA Polymerase I, Large (Klenow) Fragment (NEB), in 1x NEBuffer3. After incubation at 25 °C for 1.5 hr with gentle agitation the nuclei were washed twice with 1 ml of ice-cold TBS buffer (10 mM Tris pH 8.0, 100 mM NaCl). The nuclei pellet containing the biotin labelled DNA ends were resuspended in 500 ul ligation solution containing 0.1% Triton X-100, 50 ug Bovine Serum Albumin, and 2000 U T4 DNA Ligase (NEB) in 1x T4 DNA Ligase buffer (NEB). The sample was incubated overnight at 16 °C with gentle agitation, and the next day the nuclei were pelleted and washed with 1 ml of 1x PBS.

All the remaining steps of the protocol, reversing the formaldehyde cross-links, DNA purification and fragmentation, library preparation and Illumina sequencing were carried out as previously described ^14^. Details of the population Hi-C datasets obtained in this study can be found in Supplementary Table 1.

### Single-nucleus Hi-C protocol and samples

Single-nucleus Hi-C processing and library preparation were carried out as previously described^14,19^ using the haploid mouse ES line containing the Rex1-destabilised GFP reporter. Libraries were sequenced on an Illumina HiSeq platform with 2 x 125 bp paired end reads. Details of the single-nucleus Hi-C samples obtained in this study can be found in Supplementary Table 2.

### Hi-C sequence data processing

Both single-nucleus and population Hi-C sequencing libraries were processed using our software package “NucProcess”, which is available at: https://github.com/tjs23/nuc_processing/tree/release_1.3

Briefly, this software takes paired FASTQ sequence read files, a reference genome sequence (in this case Mouse assembly GRCm38) and knowledge of experimental parameters (restriction enzyme type and the range of size of DNA fragments sequenced in the library) to create processed, single-cell or population Hi-C contact files ^14,19^. Given that read pairs are not necessarily close in sequence, the two FASTQ files are mapped separately to the reference genome and aberrant, redundant or non-useful pairs (e.g. re-ligations) are discarded. Reads mapping to multiple sequences were kept for single-nucleus Hi-C as positional ambiguity can often be resolved and noise contacts excluded using preliminary 3D genome structures; selecting pairs that are likely close in space.

### 3D genome structure calculations

Using our previously described NucDynamics protocol^14^, 3D genome structures were calculated using simulated annealing of an initially random conformation of a particle-on-a-string representation of the chromosomes, to generate either 100 kb or 25 kb bead resolution structures that were compatible with the experimental distance restraints derived from the Hi-C contacts. Repeat calculations from different random starting positions were used to generate multiple (usually highly similar) models for each nucleus, which were then aligned spatially. The root mean square displacement (RMSD) value in particle radii between models and restraint violation data for each cell is shown in Supplementary Table 3.

The software written to perform this task and details of how to install, compile and run it (which includes both a command line version and a Jupyter notebook) is available at: https://github.com/tjs23/nuc_dynamics/tree/release_1.3

### 3D genome visualisation

Genome structures and associated data tracks were visualised using the Nuc3D software available at: https://github.com/tjs23/nuc_3d

This software reads 3D genome/chromosome structures in N3D format; a simple x,y,z coordinate format output by NucDynamics, and renders them in various styles, including as ball-and-stick, line segment or tubes. The user can then zoom, rotate, slice and expand the structures etc., and generate images and movies. Additionally, chromosome data tracks, such as those relating to contact hubs, A/B compartments or histone modifications, may be loaded (typically from a BED format file) and superimposed upon the structures, either as chromosome colours or marker symbols. We have also made a Ubuntu (Linux) virtual machine which has the Nuc3D software installed available for download at: https://zenodo.org/record/7121073#.YzSuuy1Q2Do

### ChIP-seq and ChIP-qPCR experiments

Standard protocols were used to prepare Chromatin immunoprecipitation (ChIP) samples. Briefly, fixation was carried out using 1% formaldehyde for 10 minutes at room temperature and quenched with 150 mM glycine. DNA was fragmented in the presence of 1.0% SDS using a Bioruptor sonication instrument (Diagenode) producing a size range of 200 to 300 bp. Antibodies used for ChIP are listed in Table S1, below, and the primers used for qPCR are listed in Table 2. For quantitative ChIP experiments carried out to study post-translationally modified histones, *Drosophila* spike in chromatin was added to the samples prior to the addition of the antibody. ChIP-seq libraries were prepared using the NEXTflex Rapid DNA-seq kit (Illumina) and sequenced on a Novaseq SP PE50 platform with 2 x 50 bp paired end reads at the CRUK Cambridge Institute Genomics Core facility (Cambridge, UK).

**Table S1.** Antibodies used for the ChIP-seq experiments.

| Antibody | Company | Cat no. | Host Antibody type | Amount |
| --- | --- | --- | --- | --- |
| H3K4me3 | Diagenode | pAb003-050 | rabbit polyclonal | 5 µg |
| H3K27me3 | Millipore | 07-449 | rabbit polyclonal | 5 µg |
| H3K9me3 | Active motif | 39161 | rabbit polyclonal | 5 µg |
| H3K27ac | Active motif | 39135 | rabbit polyclonal | 2.5µl |

**Table S2.**
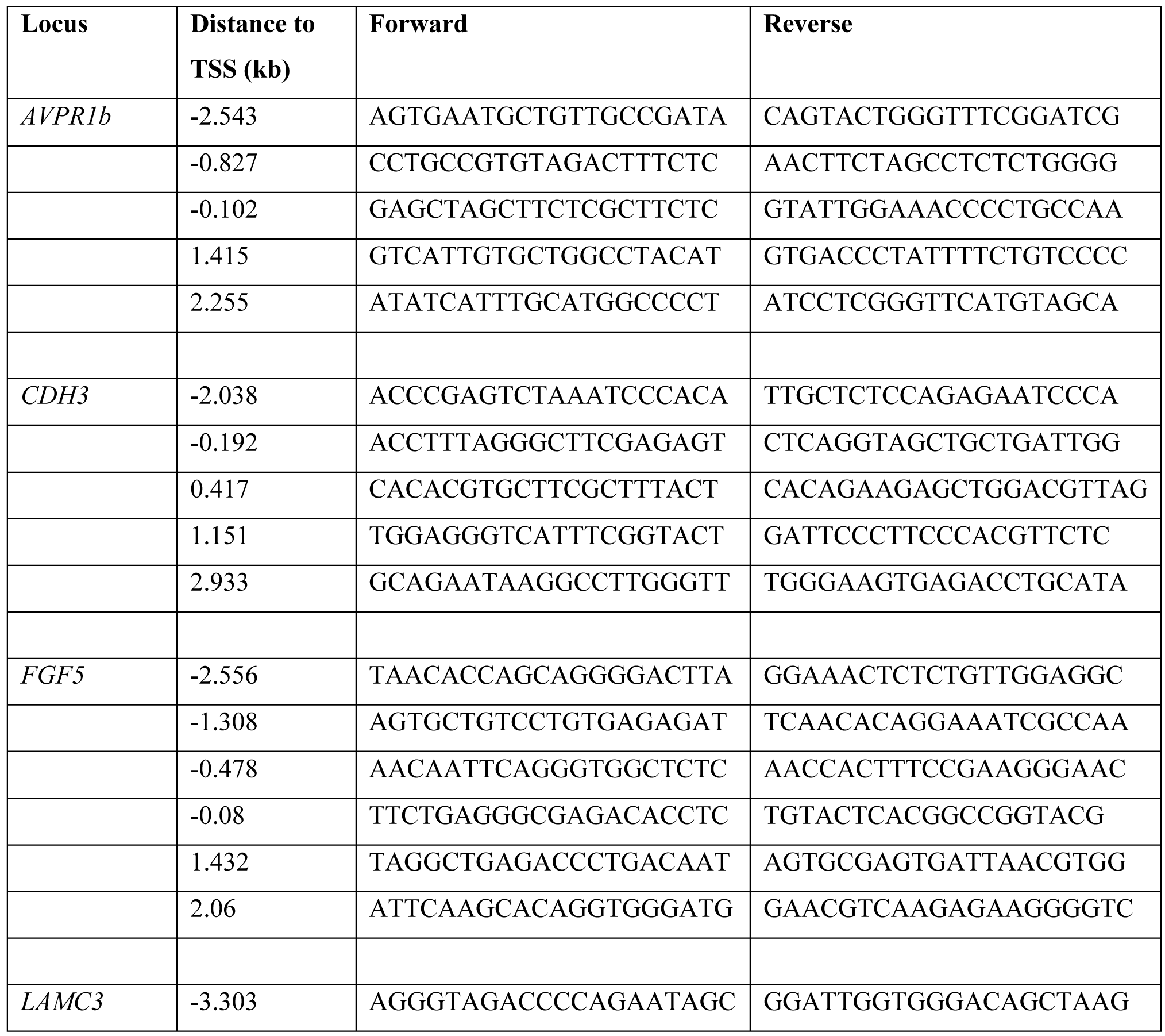
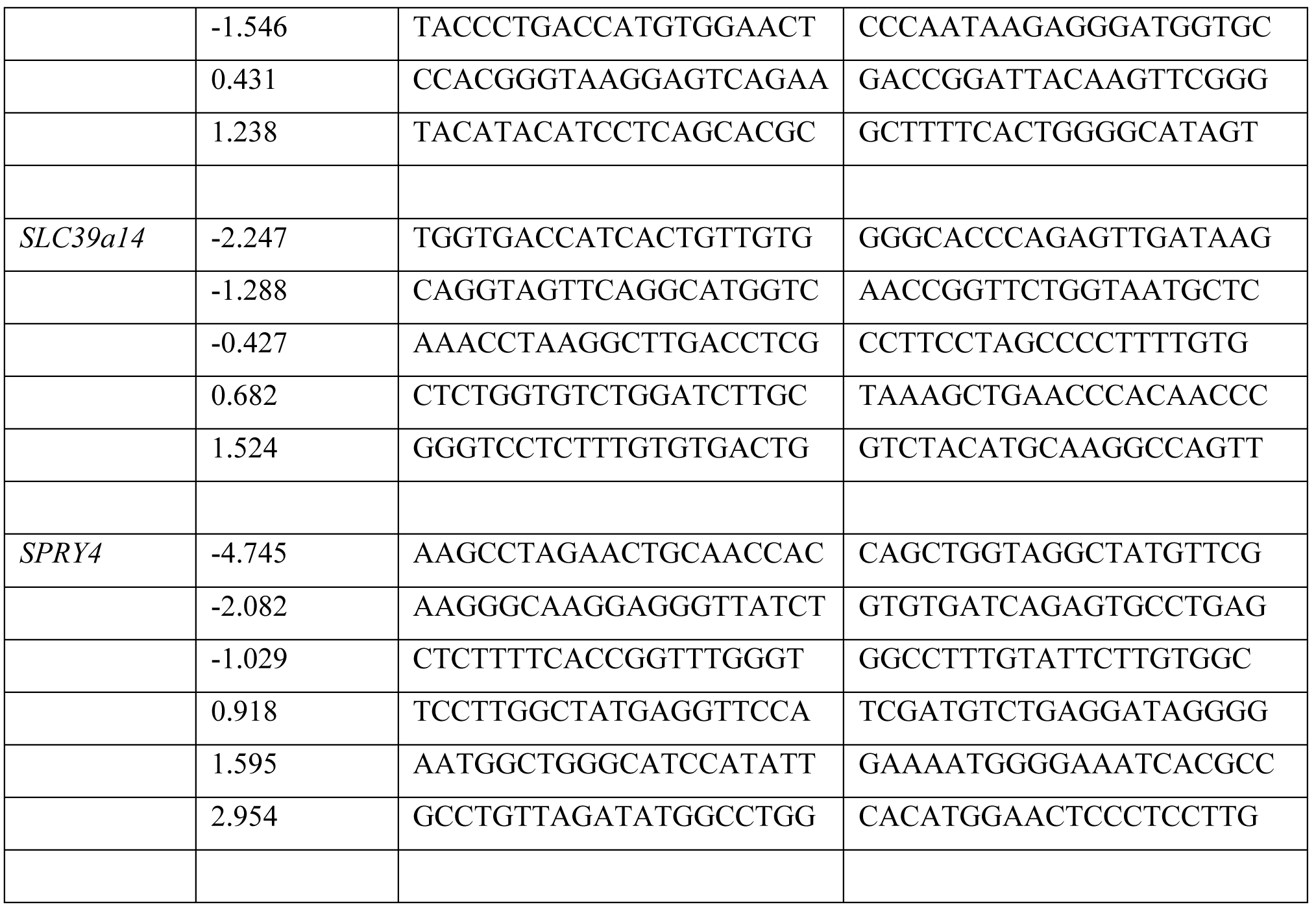
Primer sequences used for the ChIP-qPCR experiments.

### ChIP-seq data analysis

Paired-end reads were trimmed with Trim Galore v0.4.0^57^ using default parameters prior to mapping to the mouse genome reference sequence (GRCm38.p6) using Bowtie 2 v2.1.0^58^. They were then filtered to retain only those reads with a mapping quality >30 using Samtools^59^. Peaks were called using MACS2 v2.1. Model-based analysis of ChIP-Seq data was carried out using MACS2 with a minimum FDR cut-off of 0.01 (-q 0.01) to find narrowPeaks (except for H3K9me3 and H3K27me3 which have broad peak features). For samples with the *Drosophila* spike-in, reads were mapped to the *Drosophila* genome (release 6/dm6) using Bowtie 2, but with a mapping quality cut-off >10. The *Drosophila* spike-in read count was quantified using the Samtools Flagstat module. Bigwig files were normalised using the corresponding *Drosophila* spike-in read abundance and generated with the –extendReads option in deepTools^60^. ChIP-seq reads abundance profiles for each ChIP-seq peak were calculated from the normalised bigwig files using ‘bwSummary’ in Deeptools^60^.

Only peaks that appeared in both biological replicates were used (with a minimum peak overlap of 30% between replicates). To compare different time points, we generated a union set of ChIP-seq peaks for all time points and then quantified the average reads abundance of the two biological replicates at each time point.

### Defining different groups of H3K27me3 promoters

H3K27me3 ChIP-seq profile clustering at different time points was carried out using agglomerative hierachical clustering with Ward’s linkage^61^. Specifically, for each H3K27me3 bound gene promoter, ChIP-seq peak intensity differences between the 24 hr state and naïve ES cells, and between primed and 24 hr state cells, was calculated respectively for each ChIP-seq replicate. This set of intensity differences was then scaled by the maximum intensity across all time points for each gene. Euclidean distances between every pair of different promoters based on the normalised H3K27me3 intensity changes were calculated to construct a dissimilarity matrix which was used for hierarchical clustering. H3K27me3 gene clusters based on the above were defined using the ‘cutreeHybrid’ function (setting deepSplit = 1, minClusterSize = 100) within the ’dynamicTreeCut’ package in R^62^.

To understand the functional role of the different H3K27me3 promoter gene groups we analyzed genes from each cluster using the Gene ontology resource^63^ and used the Revigo tool^64^ to generate plots of the results.

### Definition of enhancer and promoter classes

The following list describes how the activity of enhancers and promoters were classified using the ChIP-seq data:

- ***Active enhancers:*** H3K27ac and no H3K4me3 or H3K27me3. (Long H3K27ac ChIP-seq peaks were trimmed to 1000 bp around the center for enhancer analysis.)
- ***Active promoters:*** Promoter TSS has H3K4me3, H3K27ac and no H3K27me3
- ***Bivalent promoters:*** Promoter TSS has H3K4me3 and H3K27me3
- ***Emerging enhancers:*** Are absent in naïve ES but present in 48 hr cells, and have at least a 3x increase in H3K27ac ChIP-seq signal when comparing 48 hr and naive ES cells
- ***Weakening enhancers:*** Are present in naïve ES but absent in 48 hr cells, and have at least a 3x decrease in H3K27ac ChIP-seq signal when comparing 48hr and naïve ES cells

### Single Cell RNA-sequencing and analysis

At each time point (Naïve ES, 24h and 48h) cells were collected using Accutase at each time point of the neuroectoderm differentiation pathway. The cell suspension was then sorted using a MoFlo cell sorter (Beckman Coulter) to recover one cell per well in a 96-well plate in 2 μl of cell lysis buffer (0.2% v/v Triton- X100 and 2U/uL of RNase inhibitor). Library preparation was carried out using the Smart-seq2 protocol^65^, and the libraries were sequenced at the CRUK Cambridge Institute Genomics Core facility (Cambridge, UK) on an Illumina HiSeq4000 sequencer.

RNA-seq libraries were aligned to the mouse reference genome (GRCm38/mm10) with the GSNAP aligner (gmap-2014-12-17)^66,67^ using the parameters -n 1 -N 1 for uniquely mapped reads and allowing for novel splice sites. Gene read counts were obtained using HTSeq (v0.6.1) based on gene annotation from Ensemble release 81. For further analysis the mapped scRNA-seq reads were filtered, normalised and analysed using the

‘Seurat v3’ package in R^68^. Specifically, low quality cells were filtered out based on a requirement for reads from a minimum of 2000 genes and 300,000 reads/cell, with less than 20% of those reads coming from mitochondrial genes. Outlier cells were further filtered using the ‘isOutlier’ function in ‘scater’^69^ (setting nmads=3, type=’lower’ and log=TRUE). Only genes with detectable expression in more than 10 cells were kept for further analysis. The most variable genes were found using the Variance Stabilizing Transform (VST) implemented in the ‘FindVariableFeatures’ function in Seurat based on a normalised expression matrix.

Potential cell cycle effects were evaluated using the ‘CellCycleScoring’ function, using previously described markers^70,71^, and cell cycle effects were then removed using the ‘ScaleData’ function (setting vars.to.regress =‘S.Score’, ‘G2M.Score’). After removal of cell cycle effects, Principle Component Analysis (PCA) of the log normalised read count for the 2000 most variable genes was carried out using the ‘RunPCA’ function in Seurat.

Pseudo-time analysis was performed using the ‘slingshot’ package in R^72^. Here PCA was carried out without scaling the genes by their expression variance using cluster labels identified by a Gaussian mixture model.

The cells were then ordered according to pseudo-time using the ‘slingPseudotime’ function. Weights assigning cells to lineages were evaluated using ‘SlingCurveWeights’. Genes with significantly variable expression over the time course were identified using the ‘tradeSeq’ package ^73^, through modelling the relationship between gene expression and pseudo-time using a Generalised Additive Model (the ‘fitGAM’ function, after obtaining a *p* < 0.01 using the ‘associationTest’ function in ‘tradeSeq’). Significantly differentially expressed genes were further classified according to their expression level changes into up-, down-, or transiently up-regulated groups through hierarchical clustering using Ward’s linkage ^61^.

### Analysis of the kinetics of gene expression

The kinetics of stochastic gene expression were investigated using the Poisson-beta statistical model^42^ and PoissonBetaSliceSampleGamma software available at: https://genomebiology.biomedcentral.com/articles/10.1186/gb-2013-14-1-r7#MOESM4

The software was run on the filtered and normalised RNA-seq reads (see above) with default settings. Genes with zero or extremely low expression were excluded, and those whose kinetics parameters could not be reliably calculated were also removed as described^42^ – in particular, this involved genes with an extremely high *k*off (i.e. >10), whose parameter estimation tends to have poor accuracy^42^.

### A/B compartment segregation

A/B compartments were defined using the Cscore tool^74^ with a bin size of 25 kb. To remove the effect of sequentially adjacent bins having higher numbers of contacts, population Hi-C reads were further normalised by dividing the Hi-C contact scores between two bins by the average Hi-C score between pairs of bins at the same sequence separation within the same chromosome. Cscores for different genomic regions were then quantile normalised and grouped according to their Cscore. For each group of genomic regions with similar Cscores (similar A/B compartment strength) we then calculated the average normalised contact scores between this group and all other Cscore groups with a different A/B compartment strength. The results were plotted in a heatmap showing the enrichment or depletion of contacts between genomic regions with different A/B compartment strength.

### Chromosome intermingling scores

Inter-chromosome intermingling scores defined as ‘the summed length of all genomic regions on other chromosomes divided by the total length of all genomic regions that are in close spatial proximity to a given locus’ were calculated from the 3D genome structures, see: https://github.com/TheLaueLab/formative-state-analysis/

To be defined as within ‘close spatial proximity’ we required the coordinates of the two genomic regions of interest to be well-defined in the 3D structure, i.e. to have an RMSD for the final 10 structural models of <1.5 bead radii. They were also required to be consistently within <2 bead radii of each other in all of the final calculated structures for a particular cell. (Varying this threshold and testing different values gave very similar results.)

To relate the intermingling scores with other features they were partitioned into 5 or 10 different groups by first quantile-normalising the scores from individual cells. For each genomic locus, we then found the median intermingling score (for that particular locus) across all the cells at a specific stage of differentiation. We then used those median scores to partition the scores into equal-sized groups (e.g. the lowest 10%, followed by 10%-to-20% etc.).

### Identification of single-cell multiway chromatin hubs and analysis of enhancer-promoter contacts

Multiway chromatin hubs were identified by searching through the single cell contact map for each chromosome with a sliding window to identify 40 kb anchor regions where a significantly greater than expected number of long-range contacts with a sequence separation of more than 50 kb are found. Where overlapping hub anchor windows were detected within a cell, we selected the one with the largest number of long-range contacts. To take account of differences in the total number of contacts and contact distribution within each cell, we estimated empirical *p*-values for the probability of finding an anchor with the observed number of long-range contacts by chance. Specifically, all long-range contacts were randomly permuted 400 times by placing the contact into a different region of the chromosome sequence keeping the sequence separation in the contact the same. We then used the same hub identification algorithm to search across each randomly permuted genome to find the number of long-range contacts to the pseudo anchor. False discovery rate (FDR) adjusted empirical *p*-values were then calculated for each hub based on the resulting distribution of the number of long-range contacts to the pseudo anchors. Only hubs with FDR adjusted empirical *p*-values of < 0.05 were kept.

To analyse the contact frequencies of regulatory elements within hubs, the different anchors and contacts were classified based on either the expression changes of genes whose TSS was < 5kb away, or the histone protein binding profile centered around the end of the contact (see ‘*Definition of enhancer and promoter classes*’, above). The distribution of the expected number of hub contacts between different regulatory elements was estimated by randomly shuffling each anchor and its group of contacts – i.e. anchors were instead paired with sets of contacts from other hubs. When performing this random shuffling, sets of contacts were selected from hubs from within the same compartment, A or B as appropriate. (This is to ensure that the contact enrichment we calculate is not simply due to a global compartmental effect.) The total number of each type of hub contact was then compared to the distribution of values from the randomly shuffled hubs – the *p* values shown are FDR adjusted empirical *p* values obtained from 5000 random permutations. The relevant scripts can be found at: https://github.com/TheLaueLab/formative-state-analysis/

### Analysis of inter-chromosomal enhancer-promoter contacts

To analyse the contact frequencies between promoters from different groups of genes and enhancers in other chromosomes, we used the single cell 3D genome structures to filter out any inter-chromosomal contacts that were not supported by the structure (i.e. not within 3.5 bead radii in the 3D polymer model). If a Hi-C contact connecting the two regulatory elements was observed within < 10 kb distance of both the gene promoter and an enhancer they were defined as being in contact. (Enhancers were classified as described in the analysis of multiway chromatin hubs, above.) The filtered single cell inter-chromosomal contacts from either all the Rex1-high or Rex1-low cells were then considered together. To obtain the distribution in the expected number of inter-chromosomal contacts between different gene promoters and enhancers, the pairing between gene promoters and enhancers was then randomly shuffled, keeping the compartment identity of both the same – i.e. genes and enhancers were randomly shuffled within either the A or B compartments, as appropriate. The total number of each type of inter-chromosomal contact was then compared to the distribution of expected values from the randomly shuffled contacts – the *p* values shown are false discovery rate (FDR) adjusted empirical *p* values obtained from 5000 random permutations.

The relevant script can be found at:https://github.com/TheLaueLab/formative-state-analysis/

---

**Supplementary Table 1.** Summary of the population Hi-C data, and statistics for the sequence analysis

| Sample | Cell type | Condition | Input reads | Mapped<br>read pairs % | Valid<br>ligation % | Total<br>contacts | Trans<br>% |
| --- | --- | --- | --- | --- | --- | --- | --- |
| Hap_ES_rep1 | Haploid | Naïve ES | 177,207,858 | 57.6 | 70.2 | 67,218,209 | 8.6 |
| Hap_ES_rep2 | Haploid | Naïve ES | 265,388,437 | 71.6 | 69.9 | 110,998,988 | 9.3 |
| Hap_24hRexH_rep1 | Haploid | 24h_Rex1-High | 52,752,503 | 56.8 | 80.4 | 23,733,171 | 40 |
| Hap_24hRexH_rep2 | Haploid | 24h_Rex1-High | 155,278,734 | 69.5 | 51.3 | 38,817,879 | 23.7 |
| Hap_24hRexL_rep1 | Haploid | 24h_Rex1-Low/formative | 65,824,401 | 57.4 | 73.9 | 27,308,454 | 40.3 |
| Hap_24hRexL_rep2 | Haploid | 24h_Rex1-Low/formative | 286,796,047 | 70.2 | 50.5 | 78,538,523 | 19.3 |
| Hap_48h_rep1 | Haploid | Primed | 172,530,619 | 50.7 | 56.1 | 34,286,170 | 24.1 |
| Hap_48h_rep2 | Haploid | Primed | 281,747,949 | 63.4 | 64.7 | 68,635,568 | 26.5 |
| RC9_ES_rep1 | RC9 diploid | Naïve ES | 43,879,391 | 70.4 | 69.0 | 18,331,335 | 11.4 |
| RC9_ES_rep2 | RC9 diploid | Naïve ES | 43,170,554 | 71.0 | 57.2 | 15,347,877 | 13.6 |
| RC9_24h_rep1 | RC9 diploid | 24h | 23,858,178 | 59.6 | 51.4 | 4,531,234 | 22.6 |
| RC9_24h_rep2 | RC9 diploid | 24h | 33,716,591 | 69.4 | 62.1 | 12,684,662 | 18.8 |
| Tet1_ES_rep1 | Tet1KO/RC9 | Naïve ES | 25,388,389 | 63.1 | 83.9 | 11,704,951 | 33.1 |
| Tet1_ES_rep2 | Tet1KO/RC9 | Naïve ES | 69,662,097 | 64.2 | 60.6 | 20,129,927 | 35.4 |
| Tet1_24h_rep1 | Tet1KO/RC9 | 24h | 28,632,985 | 64.9 | 83.9 | 13,954,500 | 35.6 |
| Tet1_24h_rep2 | Tet1KO/RC9 | 24h | 81,034,222 | 65.3 | 56.4 | 23,159,908 | 17.5 |
| Dnmt3ab_ES_rep1 | Dnmt3abKO/RC9 | Naïve ES | 20,573,708 | 63.8 | 65.3 | 8,334,738 | 11.8 |
| Dnmt3ab_ES_rep2 | Dnmt3abKO/RC9 | Naïve ES | 40,074,921 | 67.0 | 68.2 | 16,282,008 | 12.7 |
| Dnmt3ab_24h_rep1 | Dnmt3abKO/RC9 | 24h | 35,471,889 | 75.2 | 85.1 | 11,477,148 | 12.5 |
| Dnmt3ab_24h_rep2 | Dnmt3abKO/RC9 | 24h | 64,594,239 | 67.9 | 68.5 | 27,177,257 | 14.6 |

**Supplementary Table 2.** Summary of the single nucleus Hi-C data, and statistics for the sequence analysis

| Cell | GEO<br>accession | Condition | Input reads | Genome<br>mapped<br>read pairs % | valid<br>ligation % | Final<br>contact<br>pairs | Trans % |
| --- | --- | --- | --- | --- | --- | --- | --- |
| 1 | GSE80280* | Naïve ES | 1,969,076 | 62.8 | 89.4 | 110,042 | 11.7 |
| 2 | GSE80280* | Naïve ES | 1,621,648 | 58.2 | 92.3 | 65,636 | 6.3 |
| 4 | GSE80280* | Naïve ES | 1,517,614 | 46.4 | 89.9 | 75,740 | 6.4 |
| 5 | GSE80280* | Naïve ES | 1,592,161 | 55.5 | 89.8 | 61,855 | 10.1 |
| 6 | GSE80280* | Naïve ES | 1,493,430 | 43.1 | 92.2 | 60,334 | 7.7 |
| P44F6 | this study | Naïve ES | 9,631,683 | 54.9 | 85.3 | 83,849 | 11.6 |
| P44F12 | this study | Naïve ES | 8,089,512 | 54.7 | 84.9 | 53,089 | 10.5 |
| P44H4 | this study | Naïve ES | 2,496,195 | 45.0 | 82.9 | 59,919 | 7.1 |
| P62E6 | this study | 24h_Rex1-High | 8,848,740 | 56.6 | 85.2 | 110,545 | 13.2 |
| P62E12 | this study | 24h_Rex1-High | 8,167,306 | 57.5 | 79.3 | 201,451 | 9.7 |
| P62F11 | this study | 24h_Rex1-High | 9,288,453 | 54.5 | 85.5 | 111,412 | 12.1 |
| P62G7 | this study | 24h_Rex1-High | 9,909,863 | 62.8 | 50.1 | 73,969 | 15.4 |
| P62G8 | this study | 24h_Rex1-High | 11,181,667 | 54.1 | 77.1 | 171,538 | 15.2 |
| P62H10 | this study | 24h_Rex1-High | 9,512,686 | 54.9 | 81.3 | 101,843 | 14.5 |
| P62H13 | this study | 24h_Rex1-High | 10,061,265 | 53.0 | 68.9 | 170,648 | 11.8 |
| P72E10 | this study | 24h_Rex1-High | 8,852,510 | 73.2 | 38.6 | 140,169 | 15.6 |
| P63E9 | this study | 24h_Rex1-Low/formative | 13,087,790 | 55.2 | 78.3 | 153,539 | 13.3 |
| P63E14 | this study | 24h_Rex1-Low/formative | 8,999,046 | 56.3 | 69.8 | 153,526 | 9.71 |
| P63F8 | this study | 24h_Rex1-Low/formative | 8,114,855 | 50.6 | 83.6 | 89,588 | 15.6 |
| P63G10 | this study | 24h_Rex1-Low/formative | 8,902,475 | 43.6 | 73.1 | 45,180 | 15.7 |
| P63G12 | this study | 24h_Rex1-Low/formative | 8,593,456 | 56.4 | 79.2 | 138,580 | 12.1 |
| P63H7 | this study | 24h_Rex1-Low/formative | 6,663,739 | 53.7 | 79.5 | 164,059 | 9.6 |
| P63H9 | this study | 24h_Rex1-Low/formative | 6,985,362 | 56.6 | 70.0 | 32,782 | 15.4 |
| P63H10 | this study | 24h_Rex1-Low/formative | 9,830,461 | 56.4 | 81.6 | 96,883 | 15.2 |
| P63H14 | this study | 24h_Rex1-Low/formative | 5,087,395 | 30.6 | 81.7 | 76,832 | 10.4 |
| P64E5 | this study | 24h_Rex1-Low/formative | 9,132,954 | 57.3 | 62.9 | 123,707 | 14.5 |
| P64E11 | this study | 24h_Rex1-Low/formative | 8,764,583 | 59.9 | 71.6 | 170,685 | 11.3 |
| P73E8 | this study | 24h_Rex1-Low/formative | 9,729,185 | 67.2 | 47.4 | 175,663 | 23.9 |
| P73F5 | this study | 24h_Rex1-Low/formative | 9,742,947 | 48.9 | 46.9 | 171,909 | 18.2 |
| P73F6 | this study | 24h_Rex1-Low/formative | 5,638,697 | 70.9 | 47.3 | 63,916 | 22.1 |
| P73F8 | this study | 24h_Rex1-Low/formative | 6,509,055 | 75.3 | 31.4 | 90,002 | 18.6 |
| P45F10 | this study | Primed | 8,008,518 | 53.9 | 77.9 | 41,921 | 8.3 |
| P46D6 | this study | Primed | 3,088,951 | 52.9 | 75.9 | 63,035 | 6.9 |
| P46D12 | this study | Primed | 47,155,006 | 50.9 | 76.1 | 134,838 | 8.3 |
| P46G10 | this study | Primed | 7,033,068 | 53.4 | 77.9 | 47,890 | 6.2 |
| P54E14 | this study | Primed | 10,586,024 | 63.4 | 31.1 | 56,418 | 9.1 |
| P54F7 | this study | Primed | 7,276,700 | 54.8 | 77.2 | 36,819 | 7.9 |
| P54G11 | this study | Primed | 6,632,760 | 53.8 | 78.9 | 50,729 | 10.1 |
| P54G12 | this study | Primed | 7,552,400 | 49.8 | 75.8 | 60,671 | 7.4 |
| P54G13 | this study | Primed | 7,741,535 | 62.9 | 20.3 | 41,198 | 14.4 |
| P54H12 | this study | Primed | 2,669,034 | 50.0 | 81.5 | 56,298 | 10.8 |

\* Stevens, T.J., Lando, D., Basu, S., Atkinson, L.P., Cao, Y., et al. (2017), Nature 544, 59-64.

**Supplementary Table 3.**
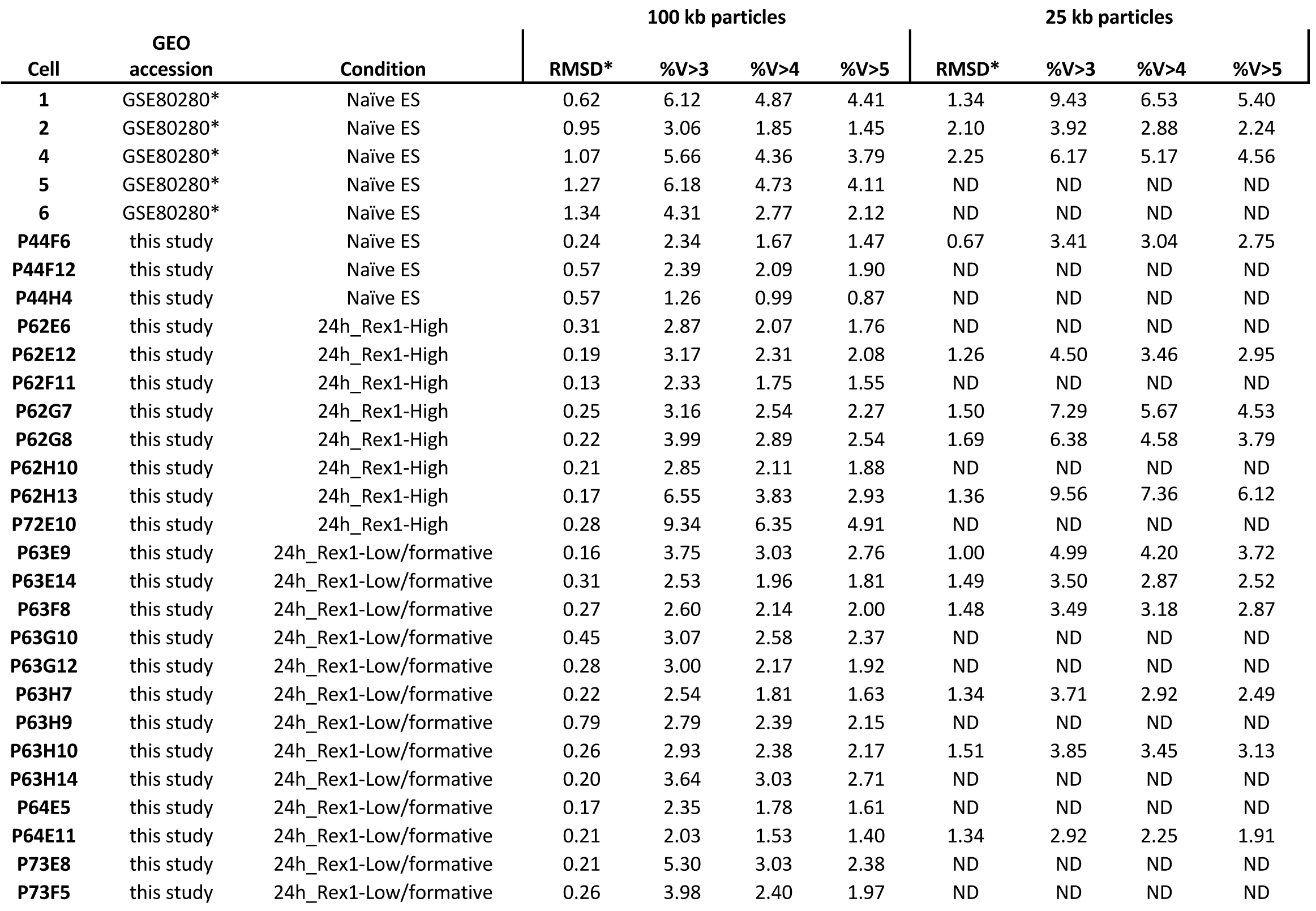
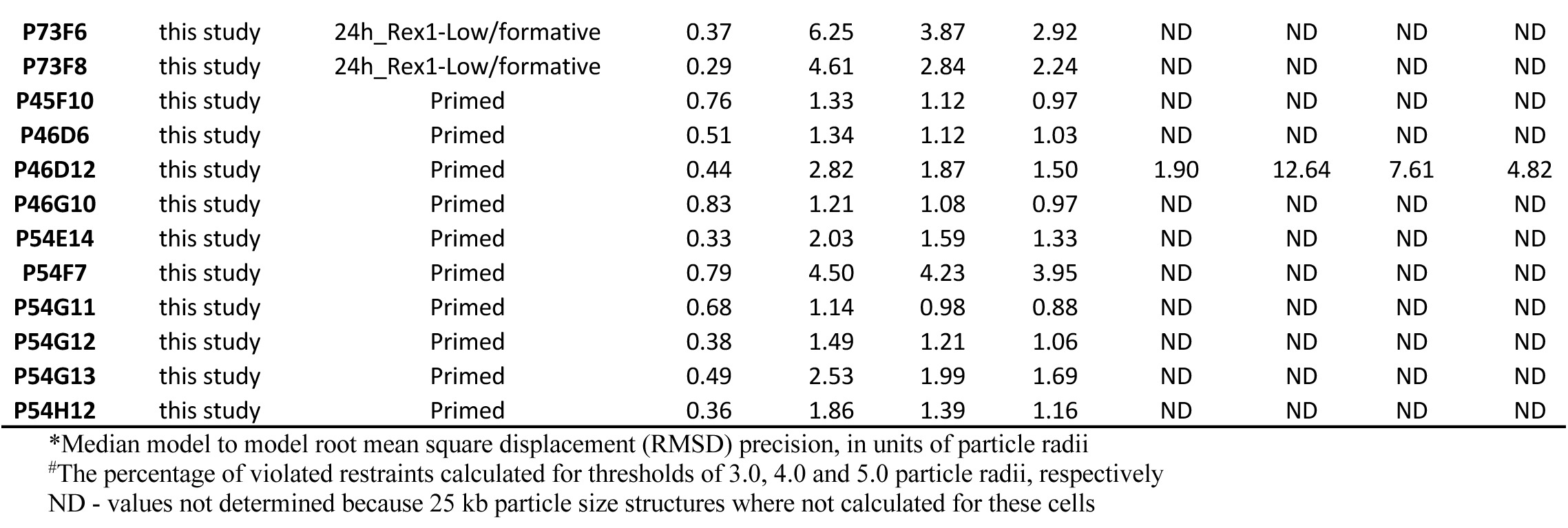
Summary of the statistics from the 3D genome structure calculation process

**Supplementary Table 4.** Classification of genes with promoters bound by H3K27 methyated nucleosomes

*(separate file)*

**Supplementary Table 5.** GO terms for Group 1 H3K27 methyated genes

*(separate file)*

**Supplementary Table 6.** GO terms for Group 2a H3K27 methyated genes

*(separate file)*

**Supplementary Table 7.** GO terms for Group 2b H3K27 methyated genes

*(separate file)*

**Supplementary Table 8.** GO terms for Group 3 H3K27 methyated genes

*(separate file)*

**Supplementary Table 9.** Classification of differentially expressed scRNA-seq genes

*(separate file)*

**Supplementary Figure 1.**
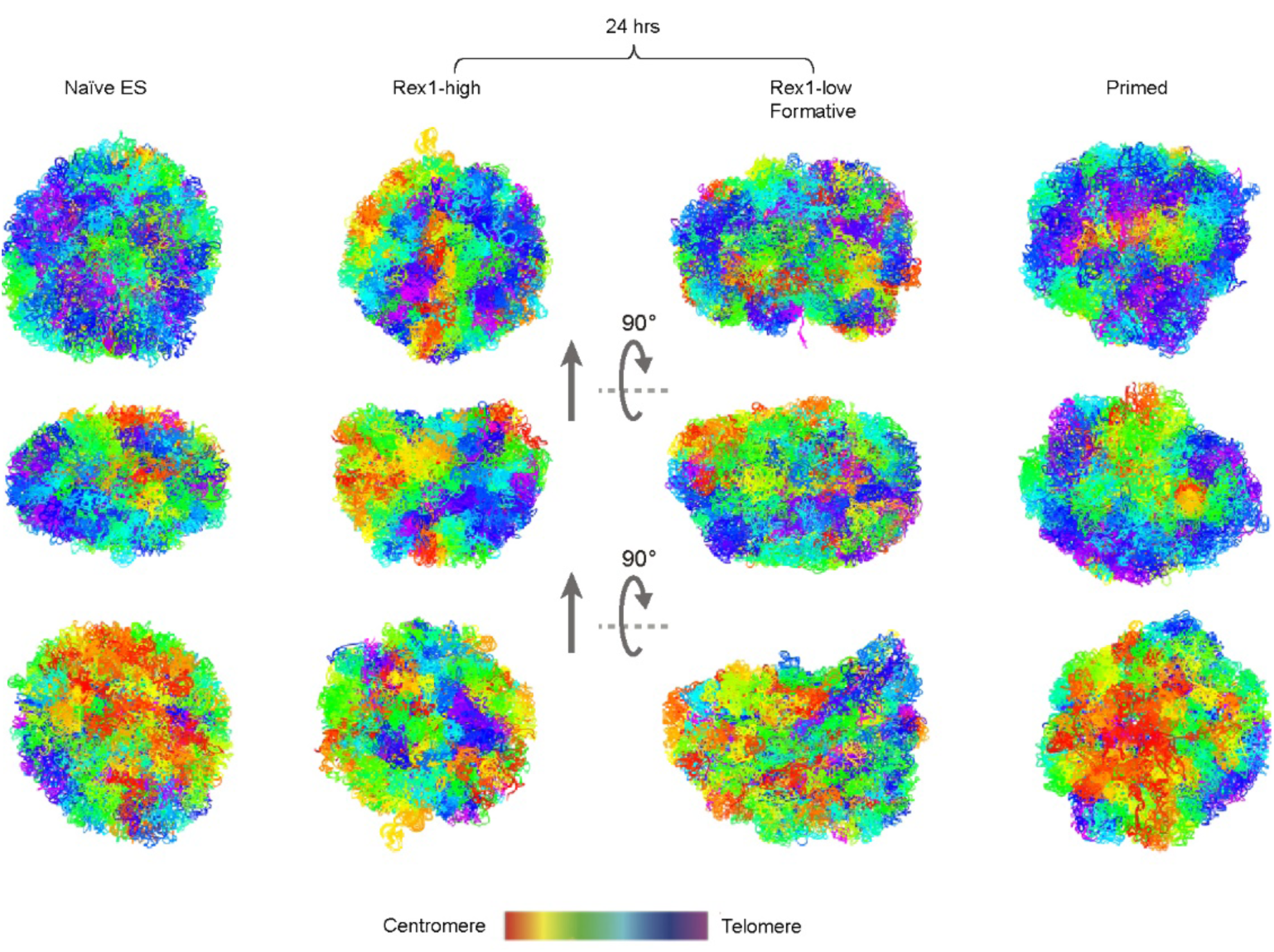
3D genome structures of representative single G1 phase nuclei, where all the chromosomes are coloured from red to blue (centromere to telomere ends), show a loss of the characteristic Rabl configuration in the 24 hr state.

**Supplementary Figure 2.**
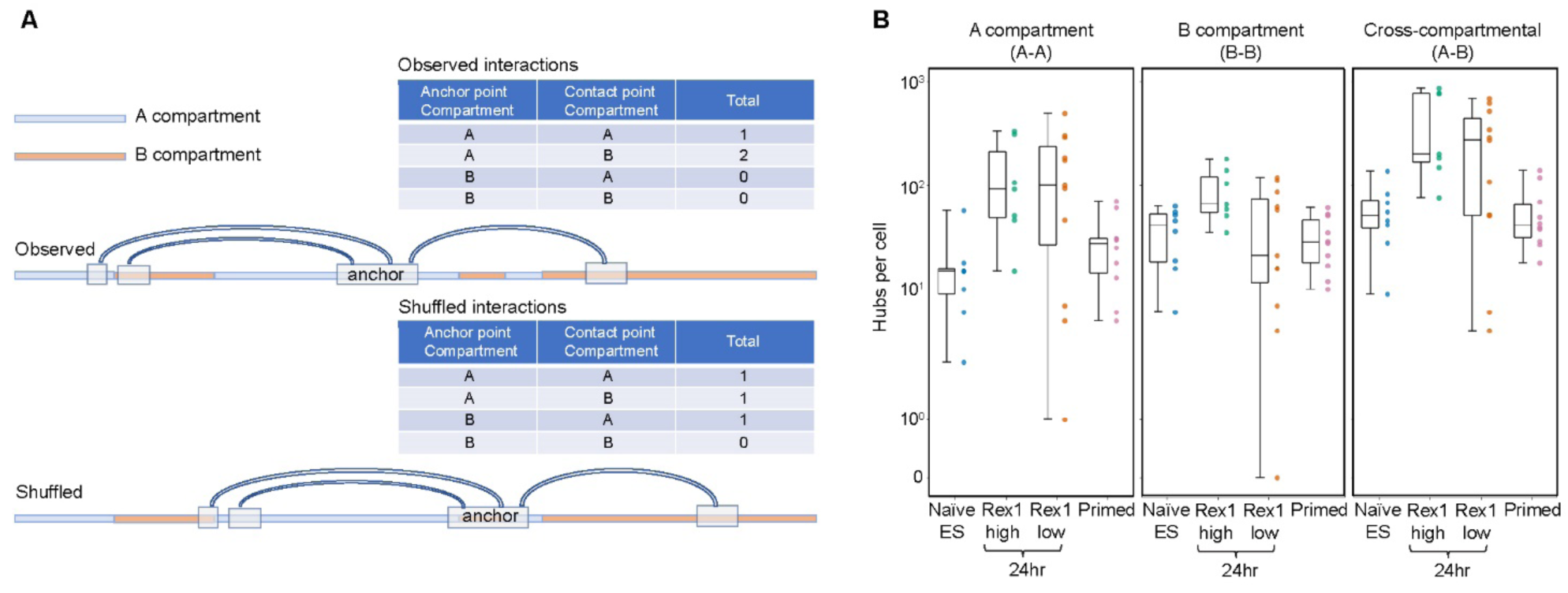
Chromatin hubs form in the A-compartment of formative state cells. **A)** The enrichment of different categories of contacts between ‘anchors’ and ‘contacts’ in chromatin hubs (compared to the random expectation) was calculated by keeping the anchors/contacts the same and shuffling the chromosomal sequence. **B)** Both Rex1-high and Rex1-low formative state cells are significantly enriched in A-compartment and A-B cross-compartmental, but not B-compartment, hubs (Mann Whitney U Test #x1D45D; = 0.0015, 0.0004 and 0.06, respectively).

**Supplementary Figure 3.**
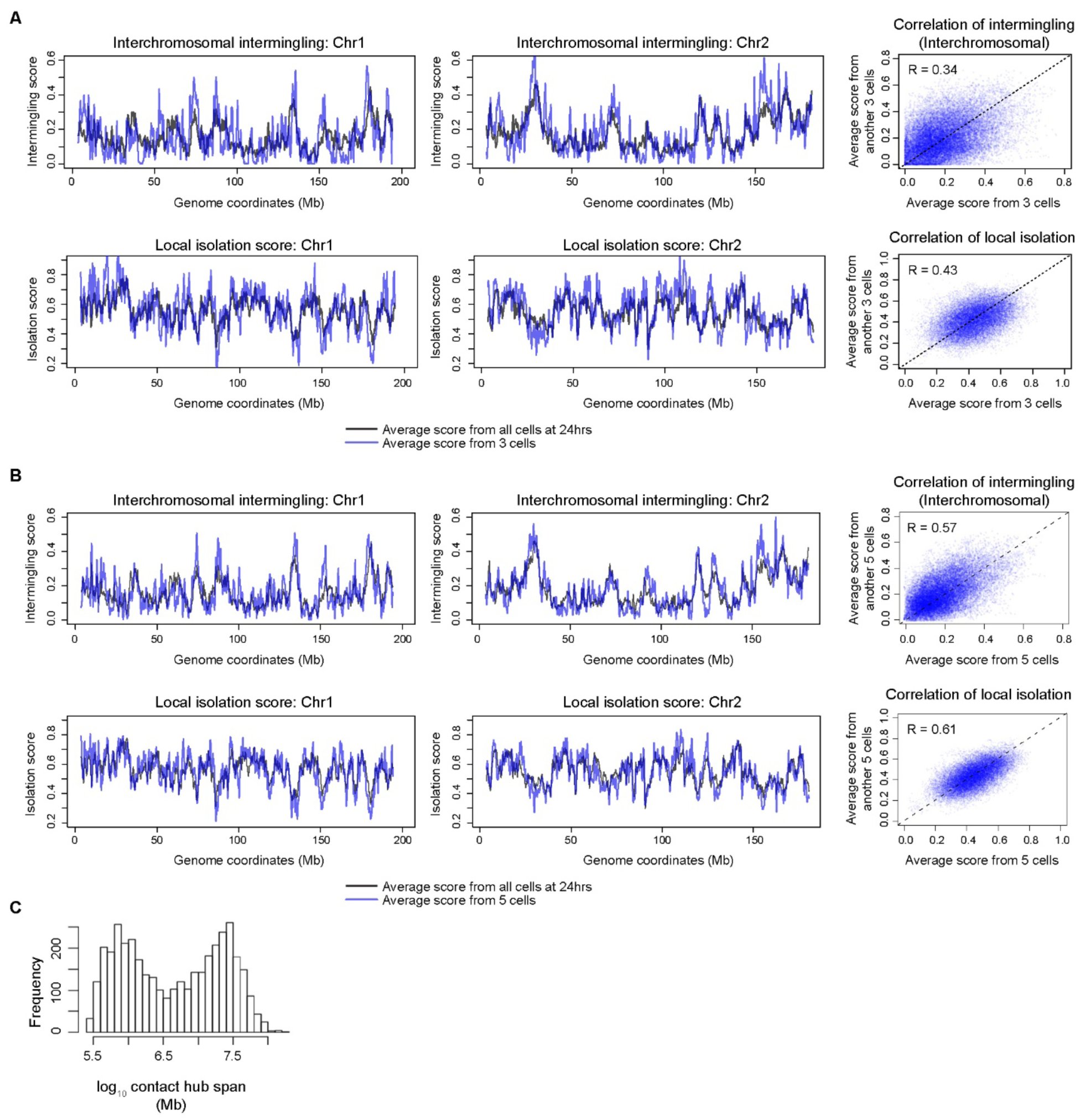
Regions of the genome that are involved in chromosomal intermingling are conserved between cells. (Left) Plots of the average intermingling scores for two representative chromosomes (1 and 2) from either three **A)** or five **B)** cells (light blue) overlaid with the average scores for the entire population of 24 hr cell structures (black). (Right) Pearson’s correlation coefficient between the average intermingling scores from either three **A)** or five **B)** cells compared to another three or five cells. (Similar results were obtained when analysing the separate Rex1-high/low cell structures – data not shown.) **C)** The histogram showing the span of genome distance for contacts in individual hubs in single cells, shows a bimodal distribution with average sizes of <1 or many Mb. (The span is defined as the log10 sequence separation between the most distant positions of intra-chromosomal contacts within a hub.)

**Supplementary Figure 4.**
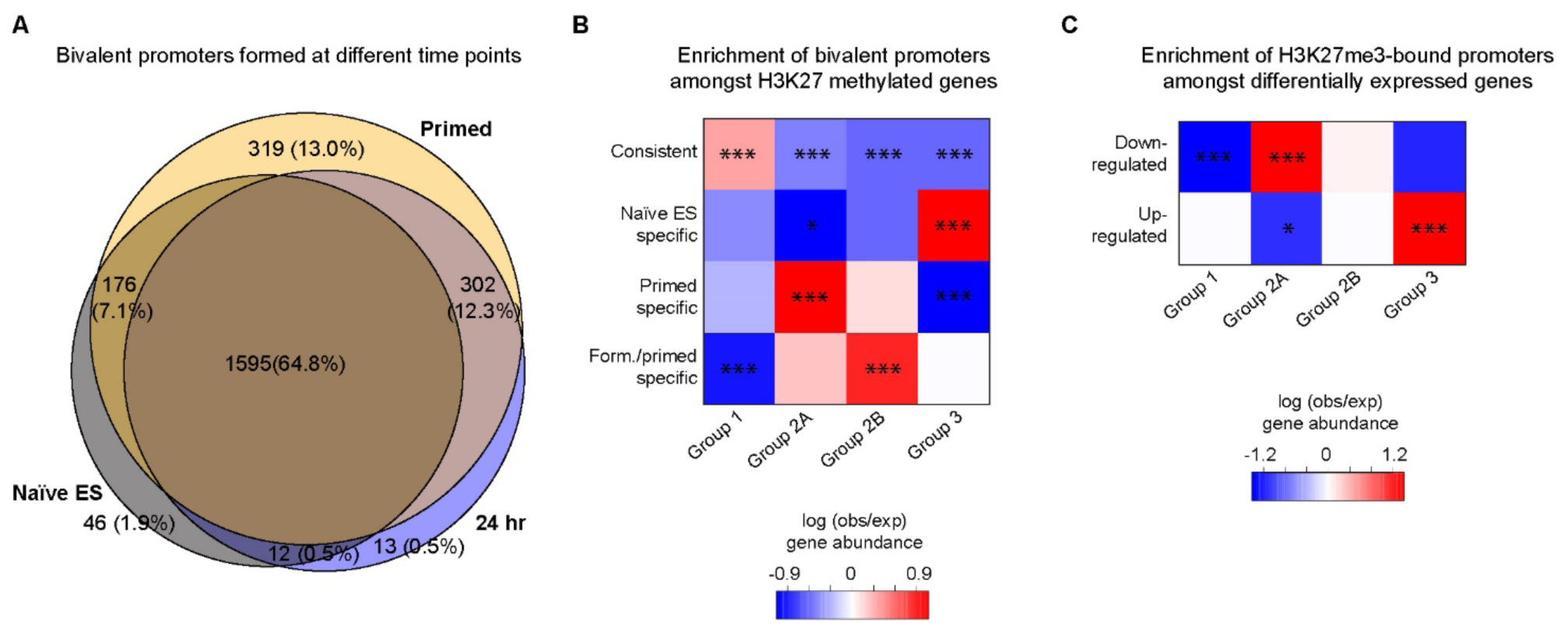
Changes in H3K4me3/H3K27me3 bivalency during the time course. **A)** The numbers of bivalent promoters (bound by H3K4me3 & H3K27me3 containing nucleosomes) formed at different time points in the differentiation pathway. **B)** The enrichment of bivalent promoters formed at different time points amongst the different groups of H3K27 methylated genes. **C)** The enrichment of H3K27me3-bound promoters amongst genes whose expression levels either increase or decrease during the time course. In B) and C) the colours indicate the observed/expected gene abundance, and FDR adjusted *p*- values (Fisher’s exact test) are shown in the heatmap where the results are significant (* - *p* < 0.05; ** - *p* < 0.01; *** - *p* < 0.001).

**Supplementary Figure 5.**
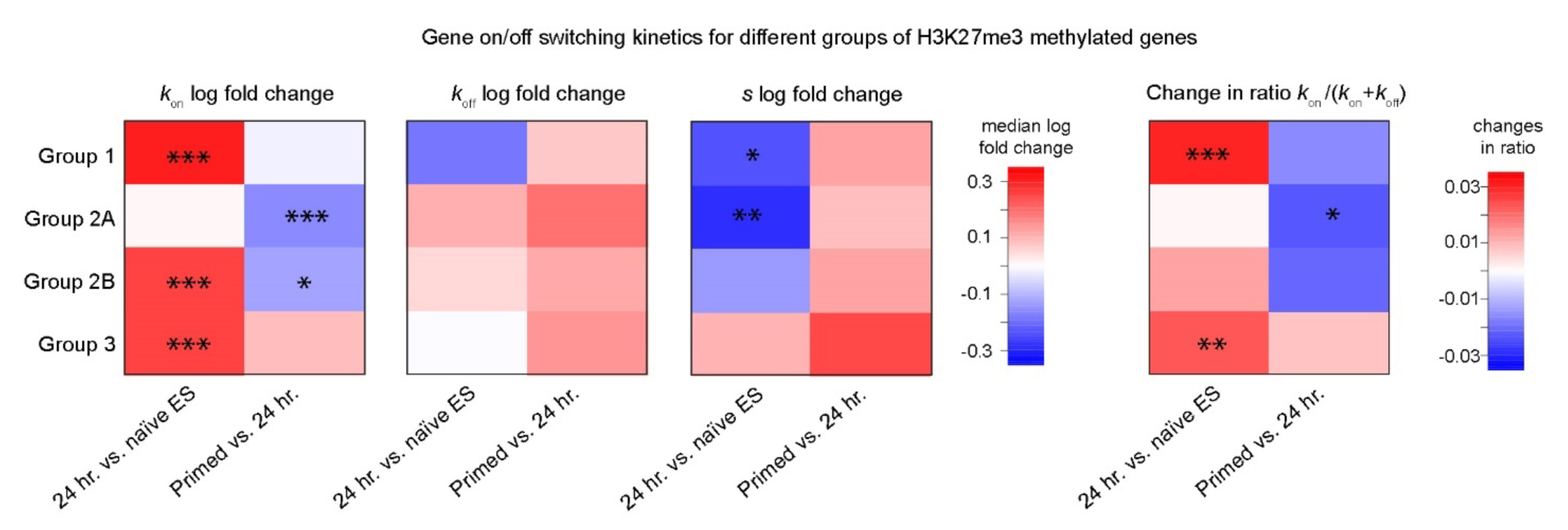
Heatmaps showing the log fold changes (Δ) in the median values of kon, koff, s and changes in the ratio of (kon/(kon+koff)) for the different groups of genes with H3K27me3-bound promoters. The colours indicate the log fold change in abundance with FDR adjusted p-values (Fisher’s exact test) shown in the heatmaps where the results are significant (* - p < 0.05; ** - p < 0.01; *** - p < 0.001).

**Supplementary Figure 6.**
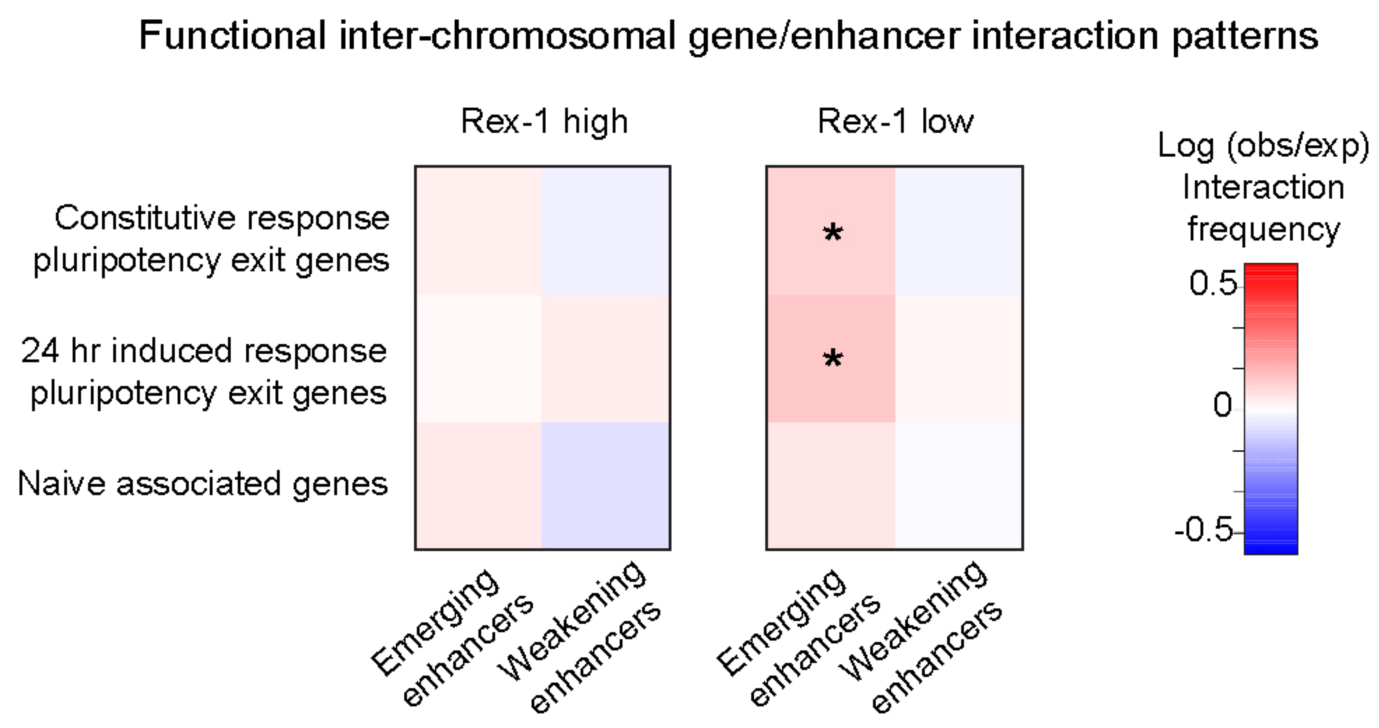
Inter-chromosomal enhancer-promoter interaction patterns in 24 hr state cells. Heatmap showing the observed vs expected interaction frequency of inter-chromosomal contacts that form within 24 hr state cells between between pluripotency exit/naïve associated promoters and emerging/weakening enhancers. The colours shown in the heatmaps indicate the log fold change in interaction frequency compared to the randomly expected distribution of the number of contacts between these regulatory elements. FDR adjusted empirical p-values are also shown (* - p < 0.05).

## Supplementary Videos

**Supplementary Video 1.** Exploded view of a naïve ES cell structure

**Supplementary Video 2.** Exploded view of a 24 hr Rex1-high structure

**Supplementary Video 3.** Exploded view of a 24 hr Rex1-low (formative state) structure.

**Supplementary Video 4.** Exploded view of a primed cell structure.

